# Hexose-6-phosphate dehydrogenase deficiency disrupts hepatic fatty acid homeostasis and induces triglyceride accumulation

**DOI:** 10.64898/2026.09.01.748570

**Authors:** Gabriele Sakalauskaite, Sophie Ebert, Julien Arthur Allard, Michael Zogg, Isabel Meister, Julia Birk, Alexander Schmidt, Jamal Bouitbir, Alex Odermatt

## Abstract

Hexose-6-phosphate dehydrogenase (H6PD) catalyzes the first two steps of an endoplasmic reticulum-specific pentose phosphate pathway, regenerating luminal NADPH levels in the process. Its function remains insufficiently well understood. Since expression of H6PD is notably high in the liver, we aimed to assess its role in hepatic metabolism. Considering the central role of the liver in lipid synthesis, breakdown and storage, we focused our efforts specifically on studying the effect of H6PD on hepatic lipid metabolism. An H6PD knockout mouse strain was generated and characterized by liquid chromatography-high-resolution mass spectrometry (LC-HRMS)-based lipidomic and proteomic analyses of liver tissue. Lipidomics analysis revealed an overall increase in hepatic triglycerides and a specific increase in unsaturated long-chain triglycerides in H6PD knockout mice. Intracellular lipid accumulation was confirmed through Nile Red staining of liver sections. Functional enrichment analysis of proteomics data from the H6PD deficient mice identified a corresponding upregulation of multiple fatty acid metabolism-associated pathways. Additionally, an H6PD knockout AML12 cell line was generated through CRISPR/Cas9 and characterized by lipid staining and functional assays to assess metabolic outcomes. Loss of H6PD led to intracellular lipid accumulation, reduced mitochondrial β-oxidation and increased sensitivity to lipotoxicity, even though fatty acids remained the cells’ primary mitochondrial fuel. Ultimately, our results indicate that H6PD plays an as-of-yet undescribed role in hepatic lipid metabolism, implying a link between the availability of NADPH within the endoplasmic reticulum and fatty acid homeostasis.

**Graphical abstract:** 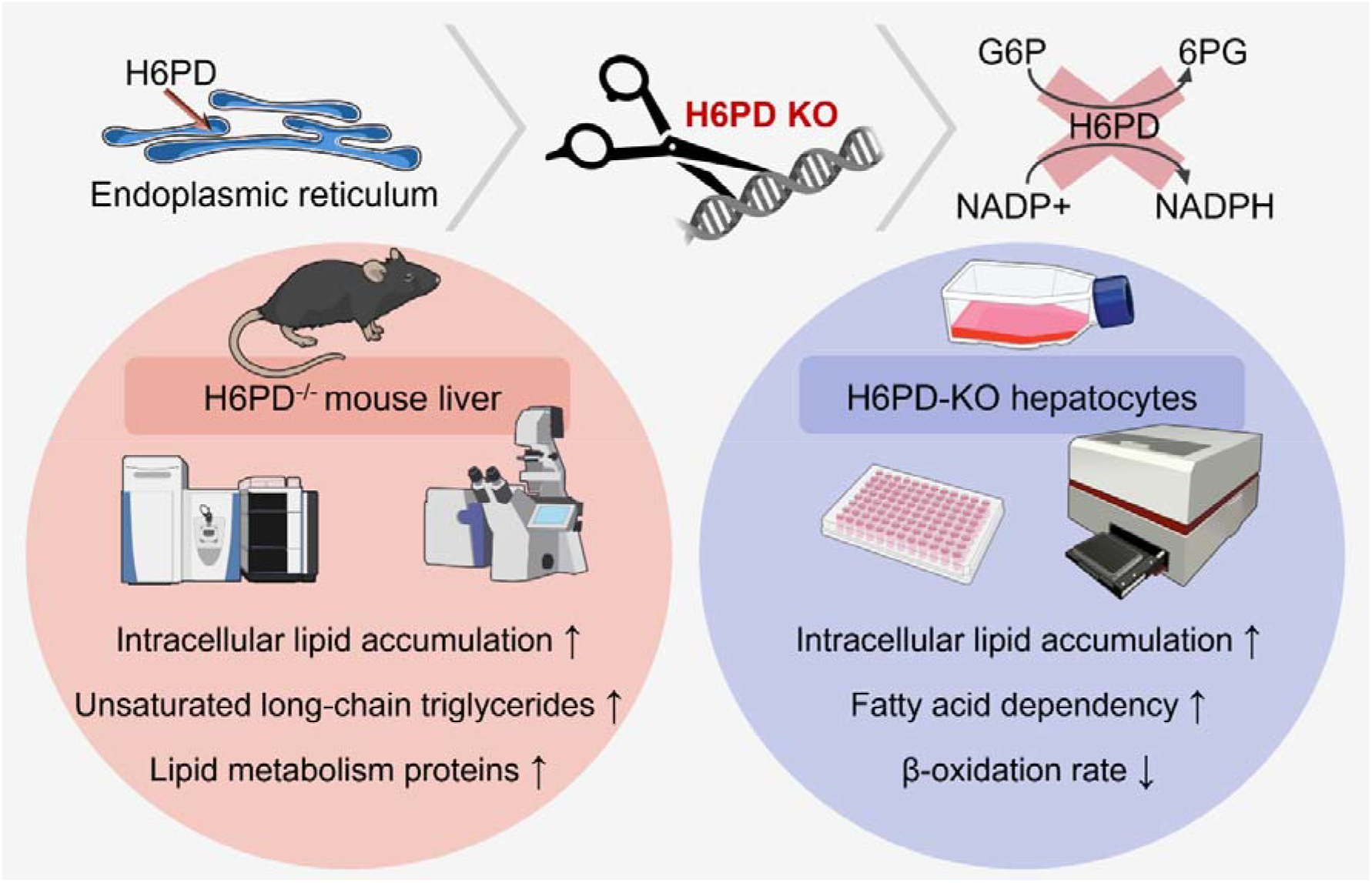

## 1. Introduction

The liver serves as a central regulator of lipid metabolism via intricate and tightly controlled pathways (1, 2). The coordinated balance of lipid synthesis, remodeling, and degradation is crucial for maintaining cellular functions, including energy storage, membrane integrity, and signaling networks (3–5). Disruption of hepatic fatty acid (FA) homeostasis impairs these regulatory mechanisms, leading to excessive triglyceride (TG) accumulation within lipid droplets, which contributes to the development and progression of metabolic dysfunction-associated steatotic liver disease (MASLD) (6, 7).

Subcellularly, the endoplasmic reticulum (ER) serves as a central hub for lipid biosynthesis and remodeling. Key metabolic processes, including the synthesis of very-long chain fatty acids (VLCFAs), TG synthesis and storage, and apolipoprotein assembly and secretion, are at least partially associated with this organelle (8–11). In response to lipotoxic stress, the ER activates adaptive mechanisms such as the unfolded protein response (UPR), alters intracellular Ca^2+^ homeostasis, and can promote lipotoxicity-induced cell death when homeostasis cannot be restored (12–14). Chronic ER stress is involved in the development of insulin resistance, type 2 diabetes and MASLD in the context of obesity (15–17).

Several enzymes involved in ER-associated lipid metabolic pathways require nicotinamide adenine dinucleotide phosphate (NADPH) as an electron donor, including 3-ketoacyl-CoA reductase (KAR) and trans-2-enoyl-CoA reductase (TECR) during VLCFA synthesis (18, 19). However, it remains unclear which of these reactions rely on the ER luminal NADPH pool. The pool is distinct from the cytosolic pool and is primarily maintained by hexose-6-phosphate dehydrogenase (H6PD) (20, 21), which catalyzes the first two reactions of an ER luminal pentose phosphate pathway (PPP)-like process, oxidizing glucose-6-phosphate to 6-phosphogluconolactone and then to 6-phosphogluconate while regenerating NADPH from NADP+ (20, 22–24). To date, the only well-studied NADPH-dependent enzyme located in the ER is 11β-hydroxysteroid dehydrogenase type 1 (11β-HSD1), which converts the inactive glucocorticoid cortisone to the stress hormone cortisol in humans (11-dehydrocorticosterone to corticosterone in rodents) (25–28). 11β-HSD1 has been shown to directly interact with H6PD (29), a ubiquitously expressed protein with particularly high expression and activity in the liver (30). H6PD knockout (KO) mouse models have provided insights into the role of this enzyme in metabolic homeostasis. Global and adipose tissue-specific H6PD KO mice exhibit fasting hypoglycemia and improved insulin sensitivity on a chow diet, indicating a role in glucose homeostasis (31, 32). Additionally, these mice exhibit impaired adipose tissue lipolysis; however, this does not translate into reduced fat mobilization or lower circulating free FA levels during fasting (32, 33). In contrast, adipose tissue-specific H6PD overexpression enhances lipolysis, while also elevating corticosterone production and modestly increasing fat mass (34). Taken together, these findings suggest that H6PD may serve as a link between ER redox homeostasis, glucocorticoid production, and carbohydrate and lipid metabolism. However, despite growing interest in its role in adipose tissue, the function of H6PD in the liver, which is a central hub for lipid metabolism, remains poorly understood.

In this study, we sought to determine whether H6PD contributes to the regulation of hepatic lipid metabolism and to characterize the effects of H6PD deficiency on FA handling and mitochondrial FA oxidation (mtFAO) in hepatocytes. To address this, we investigated the effects of H6PD loss in a mouse model as well as in the AML12 hepatocyte cell line. Lipidomics analysis, supported by lipid staining, revealed a marked accumulation of long-chain TGs with low degrees of unsaturation in the liver of H6PD-KO mice. In parallel, proteomics analysis and immunoblotting demonstrated dysregulation of proteins involved in FA metabolism, including key regulators of mtFAO. Functional analyses further showed that H6PD-deficient AML12 cells exhibited increased reliance on FAs as a fuel source, despite a reduction in mtFAO capacity, accompanied by heightened susceptibility to palmitate-induced lipotoxicity. Overall, our findings identify H6PD as an important regulator of hepatic lipid handling and mtFAO, extending its previously described roles in redox and energy metabolism to hepatic lipid homeostasis.

## 2. Materials and methods

### 2.1. Chemicals

Unless otherwise specified, all chemicals are indicated in Suppl. Table S1.

### 2.2. Creation of the H6PD-/- mice

H6PD KO mice were generated by CRISPR/Cas9 embryo electroporation. The CRISPOR software (http://crispor.tefor.net/) was used to select the target sequences aatcctgctgggagcaaccg(ggg) and agtcgctgtgatgaactcaa(ggg) in coding exon 2 of the H6PD gene. Ovulation was induced in C57BL/6J female mice by intraperitoneal injection of 5 IU equine chorionic gonadotrophin (PMSG; Folligon–InterVet, Beaucouzé, France), followed by 5 IU human chorionic gonadotropin (Pregnyl– Essex Chemie, Lucerne, Switzerland) 48 h later. Immediately after administration of human chorionic gonadotropin, the females were mated with males of the same strain. After 24 h, embryos were collected from oviducts and freed from cumulus cells by a 1–2 min treatment of 0.1% hyaluronidase (Sigma-Aldrich, Darmstadt, Germany) in M2 medium (Sigma-Aldrich). Acidic Tyrode’s solution was used to partially remove the zona pellucida. The embryos were washed and briefly cultured in M16 medium (Sigma-Aldrich) at 37°C and 5% CO_2_. Electroporation was carried out using cr:trcrRNA hybrids targeting exon 2 (8 µM each) and 16 µM Cas9 protein (all reagents from IDT, Coralville, USA) with 1 mm gap electroporation cuvettes and the ECM830 electroporator (BTX Harvard Apparatus, Holliston, USA), applying two square 3 ms pulses of 30 V with 100 ms interval as previously described (35). Of 152 electroporated embryos, 151 embryos survived. They were washed with M16 medium and transferred into the oviducts of seven 8–16-week-old pseudopregnant Crl:CD1(ICR) females. The foster mothers were allowed to deliver and raise their pups until weaning age. All produced live litters with a total of 28 viable F0 pups. 16 pups carrying the deletion of the target sequence were identified by PCR. Selected founders were bred to C57BL/6J partners to establish the H6PD KO line. The KO was confirmed by PCR (forward primer: tgattcatgccattctccaa; reverse primer: ttcccgaatgaaagccatag) and Western blot.

### 2.3. Study design and sample collection

The animal experiments were performed in agreement with the guidelines from Directive 2010/63/EU of the European Parliament on the protection of animals used for scientific purposes. All procedures were approved by the Cantonal Veterinary Office in Basel-Stadt, Switzerland (Cantonal license n° 32709_3083). The mice were housed in a standard climate-controlled facility with a 12-hour light-dark cycle and free access to standard pellet chow and water ad libitum. For our study, we included male WT and H6PD KO mice (H6PD+/+ and H6PD-/-; N = 6 per group, age 9 weeks). Mice were sacrificed by CO_2_ suffocation followed by cervical dislocation to ensure death. Blood was collected by intracardiac puncture, allowed to clot at room temperature for 30 min, centrifuged at 1,000 × g, 4°C for 10 min to separate the serum, and stored at −80°C. One piece of the liver was collected and embedded in optimal cutting temperature (OCT) compound, cooled in a pre-cooled isopentane bath in liquid nitrogen, and stored at −80°C. OCT frozen sections (10 µm) were obtained using a Cryocut 1800 (Leica Microsystems, Heerbrugg, Switzerland). The remaining part of the liver was cut into small pieces, snap-frozen, and kept at −80°C for further analysis.

### 2.4. Sample preparation for liquid chromatography coupled to LC-HRMS lipidomic analysis

Frozen liver samples of approximately 30 mg were added to tubes containing 950 mg of 1.5 mm zirconium beads and ice-cold isopropanol with 0.16 µM lysophosphatidylcholine 18:1-d7 as technical internal standard (Sigma-Aldrich, #791643C-1MG), and homogenized with a Precellys homogenizer (Bertin Technologies, Montigny le Bretonneux, France) using two cycles of 10 s at 6,000 rpm. Samples were incubated for 1 h at −20°C and centrifuged at 16,000 **×** g and 4°C for 15 min to collect two aliquots of 100 µL supernatant that were evaporated in a SpeedVac (Thermo Fisher, Waltham, MA, USA) for 1 h at low temperature setting and stored at −80°C. On the measurement day, samples were reconstituted in 150 µL of methanol, incubated for 1 h at 4°C and 900 rpm, centrifuged at 12,000 **×** g and 4°C for 15 min and transferred to LC-MS vials. QC samples were constituted by pooling 40 µL of each sample and aliquoted in three for system conditioning and injection at the start and end of the sequence. Blank samples were the resuspension solvent. The system suitability test sample was a dilute solution of UltimateSplash® (Avanti Lipids) containing 72 deuterated lipid standards.

### 2.5. LC-HRMS lipidomic measurements

Untargeted lipidomic profiles were measured on a Waters Acquity H-Class LC (Milford, MA, USA) with an Agilent Zorbax RRHD LC column (2.1 x 100 mm, 1.8 µm, Agilent) connected to a high resolution Exploris 120 orbitrap mass spectrometer (Thermo Fisher). The mobile phase A was constituted of 10 mM ammonium acetate in acetonitrile:water 6:4 (v/v) and mobile phase B of 10 mM ammonium acetate in isopropanol:acetonitrile 9:1 (v/v). The elution gradient was set at flow rate of 0.4 mL/min from 20% to 60% B in 3 min, then to 85% in 7 min and to 97% in 5 min, followed by reequilibration at the initial conditions for 4.5 column volumes. The column oven temperature was 50°C, the injection volume was 2 µL. The mass spectrometer was operated in positive mode (3.4 kV) calibrated before the start of the sequence as per manufacturer recommendations. The source settings were 50 AU for the sheath gas flow, 15 AU and 425°C for the auxiliary gas, 3 AU for the sweep gas, and the ion transfer tube temperature was set at 263°C. S-lens RF was set at 50. Each sample was acquired in profile mode with alternating full scan at 70,000 resolution on a mass range of 220-1200 m/z and top three DDA scans at 17,500 resolution with normalized collision energy of 20 eV. AGC target was set at 1e6 with an accumulation time of 100 ms. MS2 scans were triggered for signals above 1e4 at an apex of 2.5 s and dynamic exclusion of 1 s. To avoid fragmenting contaminants, a custom exclusion list based on a blank sample was used.

### 2.6. LC-HRMS lipidomic data pre-processing and data analysis

Data were converted to “.mzml” using MSConvert (36) and pre-processed in MS-DIAL version 4.7 (37). Peak detection was performed with 0.01 Da mass tolerance, minimum peak height of 200, mass slice of 0.1 Da, and smoothing using linear weighted moving average with smoothing of 3 scans and minimum peak width of 5 scans. Deconvolution of MS2 data was done with a sigma value of 0.5 and an abundance cutoff of 100. Identification relied on an embedded lipid library with a mass tolerance of 0.01 Da (MS1 and MS2) and alignment was done on the second QC sample with a retention time tolerance of 0.15 min and MS1 tolerance of 0.01 Da. Peaks in QCs with an intensity < 5x the signal observed in blanks were excluded from the peak list. Lipids were annotated with a mass tolerance of 5 ppm and spectral match. The dataset was processed in an R script, which computed CVs, applied the LOESS algorithm to correct the MS drift across the sequence and added lipid metadata. PQN correction was applied based on median QC intensities to account for variations in terms of biomass between the samples. Data analysis was done using SIMCA (Umetrics Sartorius Stedim, Umeå, Sweden) using mean centered and unit variance scaled data. Orthogonal partial least squares discriminant analysis (OPLS-DA) was performed to highlight the main trends separating H6PD+/+ and H6PD-/- in terms of lipid profiles.

### 2.7. Cell line

AML12 cells were purchased from American Type Culture Collection (ATCC, Manassas, VA, USA), tested monthly for mycoplasma, and maintained under standard conditions (37°C, 5% CO_2_). AML12 cells were cultured in Dulbecco’s Modified Eagle’s Medium/Nutrient Mixture F-12 Ham with 2.5 mM L-glutamine and 0.5 mM sodium bicarbonate (Sigma-Aldrich, #D8062) supplemented with 10% fetal bovine serum (VWR, Dietikon, Switzerland, #S00CJ1001U), 15 mM HEPES (BioConcept, Allschwil, Switzerland, #5-31F00-H), 1x ITS supplement (Sigma-Aldrich, #I3146), penicillin-streptomycin (BioConcept, #4-01F00-H) and 40 ng/mL dexamethasone (Sigma-Aldrich, #P0500-000).

### 2.8. Generation of the H6PD KO cell line

CRISPR:transfer (Cr:tr) ribonucleic acid (RNA) targeting the second exon of the murine H6PD was designed using the CHOPCHOP webtool (https://chopchop.cbu.uib.no/) (38). mH6PD crRNA (GTTACGGGCAATGTCTGCGT) was synthesized by IDT. A 50 μM cr:trRNA complex was prepared from 100 μM stock solutions of crRNA and trRNA (IDT, #1072532) with nuclease-free duplex buffer (IDT, **#**11-01-03-01) and incubated for 5 min at 95°C. Lipofection was performed using LipofectamineTM CRISPRMAX Cas9 (Thermo Fisher, #CMAX00001). cr:trRNA was mixed with Alt-R S.p. Cas9 (final concentration 1 µM each) and 25 µL Cas9plus reagent in 129 µL Opti-MEM I Reduced Serum Medium (Gibco, Reinach, Switzerland, **#**31985062) and incubated for 5 min at RT. The solution was mixed with 250 µL Opti-MEM and 15 µL CRISPRMAX transfection reagent and incubated for 20 min at RT. 500 µL transfection mix was added to a 6-well plate containing 250,000 AML12 cells in 2 mL cultivation medium. Medium was renewed after 24 h. After reaching confluence, cells were seeded at a density of 0.5 cells per well in a 96-well plate to raise single cell-derived clones. To confirm the clonality of the expanded clones, genomic DNA was extracted using the EchoLUTION Tissue DNA Micro Kit (Witec AG, Sursee, Switzerland, #010-002-250). The target site was amplified by polymerase chain reaction (PCR) (mH6PD_1_1F: ACATCGTAGCCACCTGAACC, mH6PD_1_1R: CCAGACACCAGAGTTGCTGA). PCR product was purified using the Wizard® SV Gel and PCR Clean-Up System (Promega, Dübendorf, Switzerland, **#**A9285). Following Sanger sequencing (Microsynth, Balgach, Switzerland), the TIDE webtool (https://tide.nki.nl/) was used to quantify cleavage efficiency. Protein KO was validated by Western blot (see section SDS-PAGE and immunoblotting).

### 2.9. Oleic and palmitic acid treatment

AML12 cells were seeded in 96-well plates at a density of 7,500 cells/well. A 500 mM stock solution of palmitic or oleic acid in 96% ethanol was diluted to 5 mM in sterile-filtered phosphate-buffered saline (PBS) with 5% BSA. The palmitic acid solution was heated to 37°C and the oleic acid solution to 55°C for 15 min. Treatment medium consisted of AML12 culturing medium supplemented with 75 or 150 µM BSA-conjugated oleic acid or 10, 50, and 100 µM BSA-conjugated palmitic acid. After 48 h of incubation, cells were fixed with 4% paraformaldehyde (PFA) solution and washed with PBS.

### 2.10. Nile Red and nuclei staining

Fixed AML12 cells were stained with 1x Hoechst-33342 (Life Technologies, Thermo Fisher Scientific, #H3570) and 1 µM Nile Red in PBS for 15 min, followed by three PBS washes. Frozen OCT-embedded liver sections were fixed using 4% formalin and stained with Nile Red and ROTI®Mount FluorCare DAPI (Carl Roth, Karlsruhe, Germany, #HP20.1) as described previously (39). A Cytation 5 Cell Imaging Multi-Mode Reader (BioTek, Winooski, VT, USA) with 20x objective was used to image the cells and tissue sections and measure Nile Red fluorescence intensity (469/525 nm) and cell count (377/477 nm). Total Nile Red fluorescence intensity and area were normalized to the corresponding cell count. Each experiment was carried out in six technical replicates with identical imaging settings (gain, intensity, integration time) and analysis threshold (7000). To obtain confocal images, AML12 cells were seeded on coverslips at 0.3 x 10^6^ density in 6-well plates and cultured for 48 h. Cells were fixed and stained with Nile Red and Hoechst-33258 (Invitrogen, Thermo Fisher Scientific, #H3569) as described above and mounted on microscope slides using Mowiol mounting medium. Representative images were taken using a Fluoview FV3000 confocal microscope (Olympus, Tokyo, Japan) with a 60x objective, applying consistent imaging settings for Hoechst-33258 (430/470 nm) and Nile Red (515/585 nm) across replicates.

### 2.11. TG concentration measurement

Liver tissue samples were ground to powder in liquid nitrogen using mortar and pestle. TG concentrations in mouse liver tissue and serum were measured using the Triglyceride-Glo™ Assay kit (Promega, #J3161). Luminescence was detected using the Cytation 5 reader. Results were normalized to protein concentration for all samples except serum, where equal volumes of sample were used for each replicate.

### 2.12. Preparation of microsomes

20-25 mg of liver tissue was ground to powder using liquid nitrogen. Ground tissue was resuspended in 800 µL homogenization buffer (20 mM Tris, pH 7.5, 50 mM KCl, 2 mM MgCl_2_, 0.25 M sucrose and 1x protease inhibitor cocktail) and processed by 20 strokes in a dounce homogenizer with periods of cooling on ice every five strokes for 10 s. The homogenate was centrifuged for 20 min at 4°C at 12,000 × g. The supernatant was again centrifuged for 1 h at 4°C at 104,900 × g. The microsomal pellet was resuspended in 80 µL resuspension buffer (20 mM 3-(N-morpholino)propanesulfonic acid (MOPS) pH 7.2, 100 mM KCl, 20 mM NaCl, 1 mM MgCl_2_ and 1x protease inhibitor cocktail). Protein concentration was assessed with the Pierce^TM^ BCA Protein Assay Kit (Thermo Scientific, #23225) for this assay and all the following, unless otherwise indicated.

### 2.13. Sample preparation for LC-HRMS-based proteomics analysis

The microsomal fraction was prepared as described above. Liver tissue homogenate samples were prepared by grinding 15 mg of liver tissue to powder in liquid nitrogen followed by lysis in 0.5 mL of lysis buffer (8 M urea, 75 mM NaCl, 10 mM DL-dithiothreitol (DTT), 50 mM Tris-HCl, pH 8) using strong sonication (20 cycles, 4°C, Bioruptor, Diagnode) and centrifugation at 12,000 **×** g, 10 min, 4°C. Protein concentration was determined using a reducing agent-compatible BCA assay. 25 μg of protein were combined with 25 μL 20% sodium dodecyl sulfate (SDS), 10 μL 1 M tetraethylammonium bromide (TEAB) and 5 μL 0.2 M Tris(2-carboxyethyl)phosphine hydrochloride (TCEP), the volume brought to 100 μL, and incubated at 37°C for 1 h with shaking. Iodoacetamide was added to a final concentration of 15 mM and the cysteine residues were alkylated for 30 min at 25°C in the dark. 50 μg of total protein was purified and digested with S-Trap cartridges (Protifi, Fairport NY, USA). Peptides were dried under vacuum and stored at −20°C. The samples were adjusted to a final concentration of 0.2 µg/µL with 0.1% aqueous formic acid before LC-MS/MS analysis.

### 2.14. LC-HRMS DIA proteomics measurements

0.2 µg of peptides were subjected to Liquid Chromatography with tandem mass spectrometry (LC–MS/MS) analysis using an Orbitrap Exploris 480 Mass Spectrometer fitted with a Vanquish Neo (both Thermo Fisher Scientific) and a custom-made column heater set to 60°C. Peptides were resolved using a RP-HPLC column (75 μm × 30 cm) packed in-house with C18 resin (ReproSil-Pur C18–AQ, 1.9 μm resin; Dr. Maisch GmbH) at a flow rate of 0.2 μL/min. Separation of peptides was achieved using the following gradient: 4% Buffer B to 10% Buffer B in 5 min, 10% Buffer B to 35% Buffer B in 45 min, 35% Buffer B to 50% Buffer B in 10 min. Buffer A was 0.1% formic acid in water and Buffer B was 80% acetonitrile, 0.1% formic acid in water. The mass spectrometer was operated in DIA acquisition mode with a total cycle time not exceeding approximately 3 s. For MS1, the following parameters were set: resolution: 120,000 FWHM (at 200 m/z), scan range: 350-1400 m/z, injection time: 25 ms, normalized AGC target: 300%. MS2 (SWATH) scans were acquired using the following parameters: isolation window: 8 m/z, HCD collision energy (normalized): 28%, normalized AGC target: 1000%, resolution: 15,000 FWHM (at 200 m/z), precursor mass range: 400 – 900 m/z, max. fill time: 22 ms, data type: centroid. In total 100 DIA (MS2) mass windows followed one MS1 scan per MS cycle.

### 2.15. Proteomics data analysis

The acquired data were searched using SpectroNaut (v16.0, default settings) against a human/mouse database (consisting of 20360/17085 protein sequences downloaded from Uniprot on 2022-02-22) and 392 commonly observed contaminants using the following search criteria: full tryptic specificity was required (cleavage after lysine or arginine residues, unless followed by proline); three missed cleavages were allowed; carbamidomethylation (C) was set as fixed modification; oxidation (M), N-acetylation (N-term) were applied as variable modifications. The raw quantitative data was further statistically analyzed using MSstats (40). All proteins detected by at least 2 unique peptides were ranked by sign(log2FC)*(-log10(pvalue)) and subjected to gene set enrichment analysis (GSEA) using clusterProfiler v4.18.4 (41) in R v4.5.1 (42) with the Hallmark gene set provided by the Molecular Signatures Database. The 300 most significantly altered proteins were additionally subjected to functional enrichment analysis using the Kyoto Encyclopedia of Genes and Genomes (KEGG) pathway database within STRING (43). Results were ranked by strength (log10(observed proteins/expected proteins)) and false discovery rate (FDR) shown as p-value corrected for multiple testing using the Benjamini-Hochberg procedure.

### 2.16. Seahorse analysis

Oxygen consumption rate (OCR) and extracellular acidification rate (ECAR) were determined in AML12 cells using a Seahorse XFp Extracellular Flux Analyzer (Agilent Technologies, Santa Clara, CA, USA). 5,000 cells/well were seeded in XFp plates and incubated for 24 h. Cells were washed three times with prewarmed XF assay modified DMEM medium (Agilent Technologies, #103681-100) supplemented with 10 mM glucose (Agilent Technologies, #103577-100), 2 mM glutamine (Agilent Technologies, #103578-100) and 1 mM pyruvate (Agilent Technologies, #103579-100) and incubated for one hour at 37°C in a non-CO_2_ incubator. Mito Fuel Flex Test was performed according to the manufacturer’s protocol. OCR was measured at baseline and following injections of UK5099 (Sigma-Aldrich, #5048170001, 2 μM), BPTES (Sigma-Aldrich, #SML0601, 6 μM), and etomoxir (Sigma-Aldrich, #236020, 8 μM), inhibiting the mitochondrial import of the three main metabolic substrates pyruvate, glutamine, and fatty acids, respectively. The experiment was performed in technical triplicates three times independently. OCR and ECAR were automatically calculated by Seahorse XFp software version 2.2.0 (Seahorse Bioscience, Billerica, MA, USA) and normalized to protein content.

### 2.17. Fatty acid uptake assay

AML12 cells were seeded at 15,000 cells/well density in 96-well-plates and incubated in regular medium for 24 h. Cells were washed with PBS and starved for 30 min in phenol red-free DMEM/F-12 Ham (Sigma-Aldrich, #D6434) without supplements. Cells were then treated with 1 µM green fluorescent PA (BODIPY™ FL C16, Thermo Fisher Scientific, #D3821) in phenol-red free DMEM/F-12 Ham. Cells were fixed using 4% PFA after 30, 45, 60, 90, 120, and 150 min and stained with Hoechst-33258 as described before. Mean fluorescence intensities of the green fluorescent PA and Hoechst-33258 were measured using the Cytation 5 reader with excitation/emission wavelengths of 488/530 nm and 400/439 nm, respectively. Fluorescence intensity of green fluorescent PA was normalized to Hoechst-33258 fluorescence intensity. Three biological replicates with three technical replicates each were performed.

### 2.18. β-oxidation assay using radioactively labeled PA

AML12 cells were seeded at 1 x 10^4^ cells/well density in regular medium in 96-well plates. After 24 h, medium was exchanged to fresh regular medium or treatment medium, supplemented with 10 or 100 µM etomoxir (Sigma-Aldrich, #236020). Cells were incubated for 2 h, washed with PBS and incubated for 3 h in radioactive assay medium (DMEM/HAM F12 phenol red-free medium supplemented with 10 mg/mL FA free bovine serum albumin (BSA) (Sigma-Aldrich, #A8806), 100 µM PA, 1 mM L-carnitine (Sigma-Aldrich, #C0283) and [1-^14^C] PA (Revvity, Waltham, MA, USA, #NEC075H050UC) at a 1:500 dilution). Cells were precipitated using 6% perchloric acid at 37°C for 10 min. The supernatant was centrifuged for 10 min at 2,000 **×** g and transferred into vials containing 3 mL Ultima Gold TM XR scintillation cocktail (Revvity, #6013117). Aliquots of assay medium with and without [1-^14^C] PA were used as positive and negative controls. Sample radioactivity was measured with a Tri-Carb scintillation counter (Revvity) using a count time of 2 min. A corresponding plate, treated for 3 h with assay medium without [1-^14^C] PA, was fixed using 4% PFA and stained with Hoechst-33258. Nuclei were counted using the Cytation 5 reader. Radioactivity measurements were normalized to averaged cell numbers. Four technical replicates were performed per experiment, with four biological replicates in total. Outliers of technical replicates were excluded after outlier analysis (GraphPad, α = 0.05), a minimum of two technical replicates were used for quantification.

### 2.19. SDS-PAGE and immunoblotting

1 x 10^6^ AML12 cells were seeded in 10 cm culture plates and cultivated for 48 h. Cells were washed twice with PBS and lysed with 1 mL radioimmunoprecipitation assay (RIPA) buffer containing 1x protease inhibitor and 1x phosphatase inhibitor. For tissue samples, 20 mg of mouse liver tissue were homogenized with Precellys 1.4 mm zirconium oxide beads (Bertin Technologies, #P000927-LYSKO-A) in 1 mL lysis buffer with a Precellys® Evolution Touch tissue homogenizer (Bertin Technologies). Sodium dodecyl sulfate polyacrylamide gel electrophoresis (SDS-PAGE) and immunoblotting were performed as previously described (44). Primary and secondary antibodies with corresponding sample reducing agent, antibody diluent and blocking solution are listed in Suppl. Table S2. 15-30 µg of protein were used for SDS-PAGE. Precision Plus Protein All Blue Standards (BioRad Laboratories AG, Cressier, Switzerland, #1610373) served as protein ladder. Proteins were transferred to Polyvinylidene fluoride membranes with a pore size of 0.45 µm (Millipore, Sigma-Aldrich, #IPVH00010) using Trans-Blot® SD Semi-Dry Transfer (Bio-Rad) and detected using Immobilon® Western Chemiluminescent HRP Substrate (Millipore, Sigma-Aldrich #WBKLS0500). Densitometry analysis was carried out using ImageJ software (version 1.53i, RRID:SCR_003070). Protein band intensities were normalized to the mean Ponceau staining intensity of the corresponding lane, following background signal subtraction.

### 2.20 RNA isolation and RT-qPCR

RNA was isolated from AML12 cells using the RNeasy Mini Kit (Qiagen, Venlo, Netherlands, # 74106). Reverse transcription (RT) was performed using the PrimeScript RT Reagent Kit (Takara, Kyoto, Japan, #RR037BA) for 25 min at 37°C and 5 s at 85°C. Real-Time Quantitative PCR (RT-qPCR) was conducted in technical triplicates using 5 ng cDNA per reaction and the KAPA SYBR FAST qPCR Kit (Roche, Sigma-Aldrich, #KK4619) with the QuantStudio 6 Pro Real-Time PCR System (Applied Biosystems, Thermo Fisher Scientific). Primer sequences and the run method are provided in Suppl. Table S3. Data were analyzed using QuantStudio Design and Analysis Software 2.8.0.

### 2.21. Statistical analysis

GraphPad Prism software 8.0 (GraphPad, La Jolla, CA, USA, RRID:SCR_002798) and Microsoft Excel (Microsoft, Redmond, WA, USA) were used for data analysis unless otherwise indicated. Ordinary two-way analysis of variance (ANOVA) followed by Fisher’s Least Significant Difference (LSD) post hoc test was performed when comparing multiple groups. An unpaired t test with Welch’s correction was performed when comparing two groups. A p-value < 0.05 was considered statistically significant. Bar graphs indicate mean ± standard deviation (SD). Asterisks without brackets indicate comparison to the respective control treatment.

## 3. Results

### 3.1. Levels of unsaturated long-chain TGs are elevated in the livers of H6PD-deficient mice

The *H6pd* gene was disrupted in mice using CRISPR/Cas9-mediated genome editing, and successful KO was confirmed by Western blot analysis of liver tissue (Suppl. Fig. S1). To characterize the metabolic consequences of H6PD deficiency *in vivo*, we performed a comprehensive lipidomic analysis using our validated global H6PD KO mouse model. Hepatic lipid profiles from H6PD+/+ and H6PD-/- mice were analyzed by LC-HRMS-based lipidomics (Fig. 1). Lipid class analysis revealed that phosphatidylcholines (PC) and lysophosphatidylethanolamines (LPE) were among the most significantly decreased lipid classes in H6PD-/- livers compared with H6PD+/+ controls (Fig. 1A). Additionally, levels of lysophosphatidylcholines (LPC), a major product of PC hydrolysis, were also reduced in H6PD-/- mice. Together, these changes suggest altered phospholipid metabolism and impaired phospholipid remodeling. Conversely, TGs and oxidized triacylglycerols (oxTGs) were significantly elevated in livers of H6PD-/- mice (Fig. 1A).

**Figure 1.**
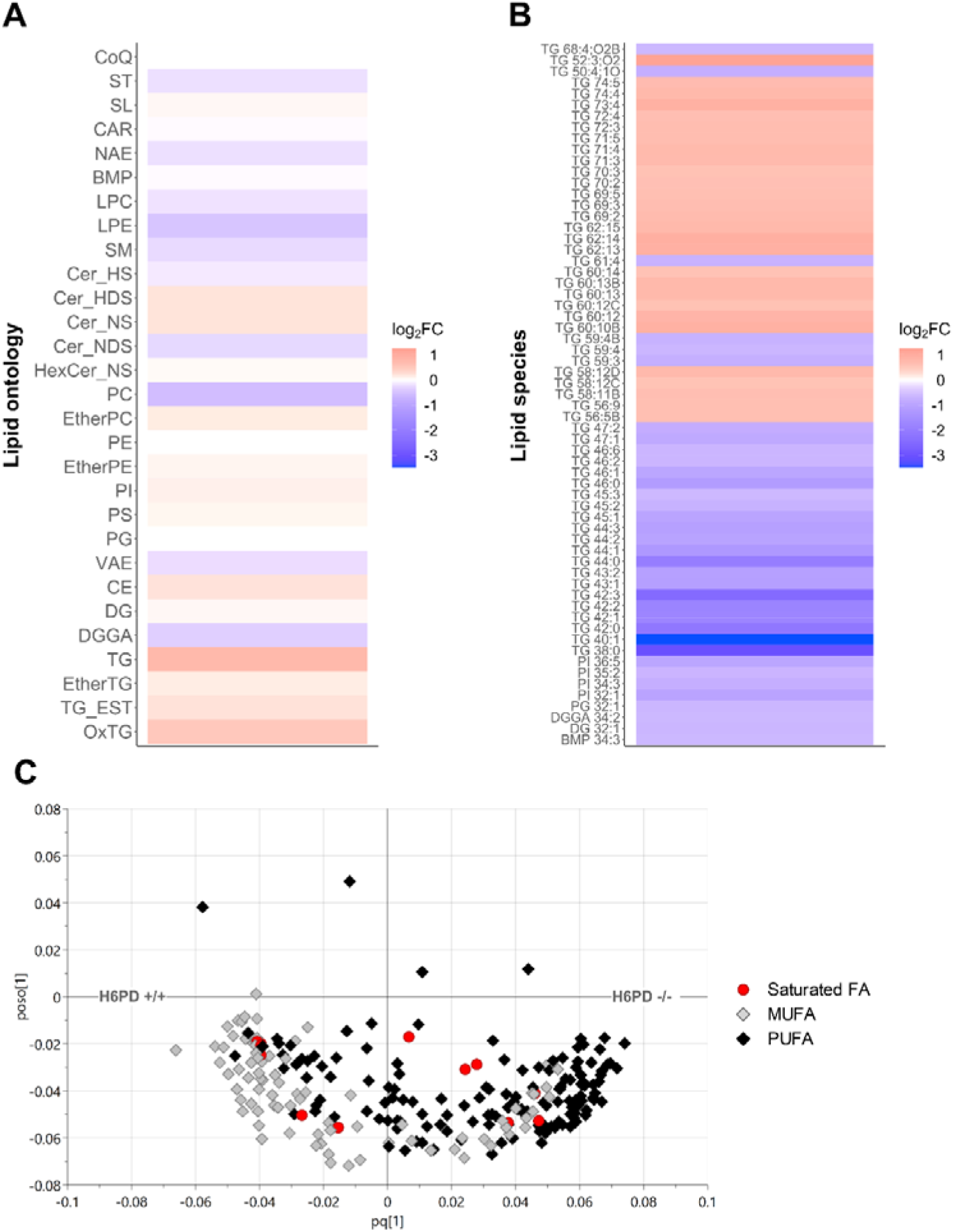
Lipidomic analysis reveals altered hepatic lipid composition in H6PD-deficient mice. (**A**, **B**) Heatmaps showing the log_2_ fold change in hepatic lipid abundance in H6PD-/- mice relative to H6PD+/+ controls. (**A**) Fold changes in lipid abundance summed across each indicated lipid class. (**B**) Heatmap of individual lipid species exhibiting the largest changes in abundance (|log_2_ FC| > 1.5), arranged by ontology and retention time. (**C**) OPLS-DA loadings plot highlighting TG species contributing to the separation between H6PD+/+ and H6PD-/- (model with one predictive and two orthogonal components, R^2^X = 0.64; Q^2^ = 0.63, variables = 648). Symbols indicate the degree of fatty acyl unsaturation (red circle, saturated; grey diamond, monounsaturated fatty acid (MUFA) with 1 double bond; black diamond, polyunsaturated fatty acid (PUFA) with ≥ 2 double bonds).

To further characterize these alterations, TG species were classified according to fatty acyl chain length and degree of unsaturation. This analysis revealed a marked reduction in short- and medium-chain TG species, accompanied by an accumulation of long-chain TGs (Fig. 1B). Further stratification by the number of double bonds demonstrated that this increase was driven predominantly by highly unsaturated TG species (Fig. 1B, C). Overall, these findings indicate that H6PD deficiency profoundly remodels the hepatic lipidome, shifting TG composition toward longer-chain, more highly unsaturated TG species, while simultaneously disrupting phospholipid homeostasis.

### 3.2. H6PD deficiency in mouse liver and AML12 cells leads to increased intracellular lipid storage

To validate the hepatic lipid accumulation observed in the lipidomic analysis, liver sections from H6PD+/+ and H6PD-/- mice were co-stained with Hoechst and the neutral lipid dye Nile Red. Confocal microscopy revealed a marked increase in Nile Red-positive lipid droplets in H6PD-/- livers compared with H6PD+/+ controls (Suppl. Fig. S2A). To quantify this accumulation, Nile Red-stained liver sections were imaged using a Cytation 5 high content imaging system with automated image analysis (Fig. 2A, B). Quantification demonstrated significantly higher Nile Red fluorescence intensity in H6PD-/- livers than in H6PD+/+ controls (Fig. 2B). Consistent with this finding, a luminescence-based assay detected a trend towards increased hepatic TG content in H6PD-/- mice (Suppl. Fig. S2B). In contrast, serum TG concentrations, as well as hepatic and serum free glycerol levels, were not significantly different between genotypes (Suppl. Fig. S2B, C, respectively). Collectively, these findings indicate that H6PD deficiency promotes hepatic neutral lipid accumulation without detectable alterations in hepatic or circulating glycerol levels, suggesting that the increased hepatic TG content is associated with altered intracellular lipid storage.

**Figure 2.**
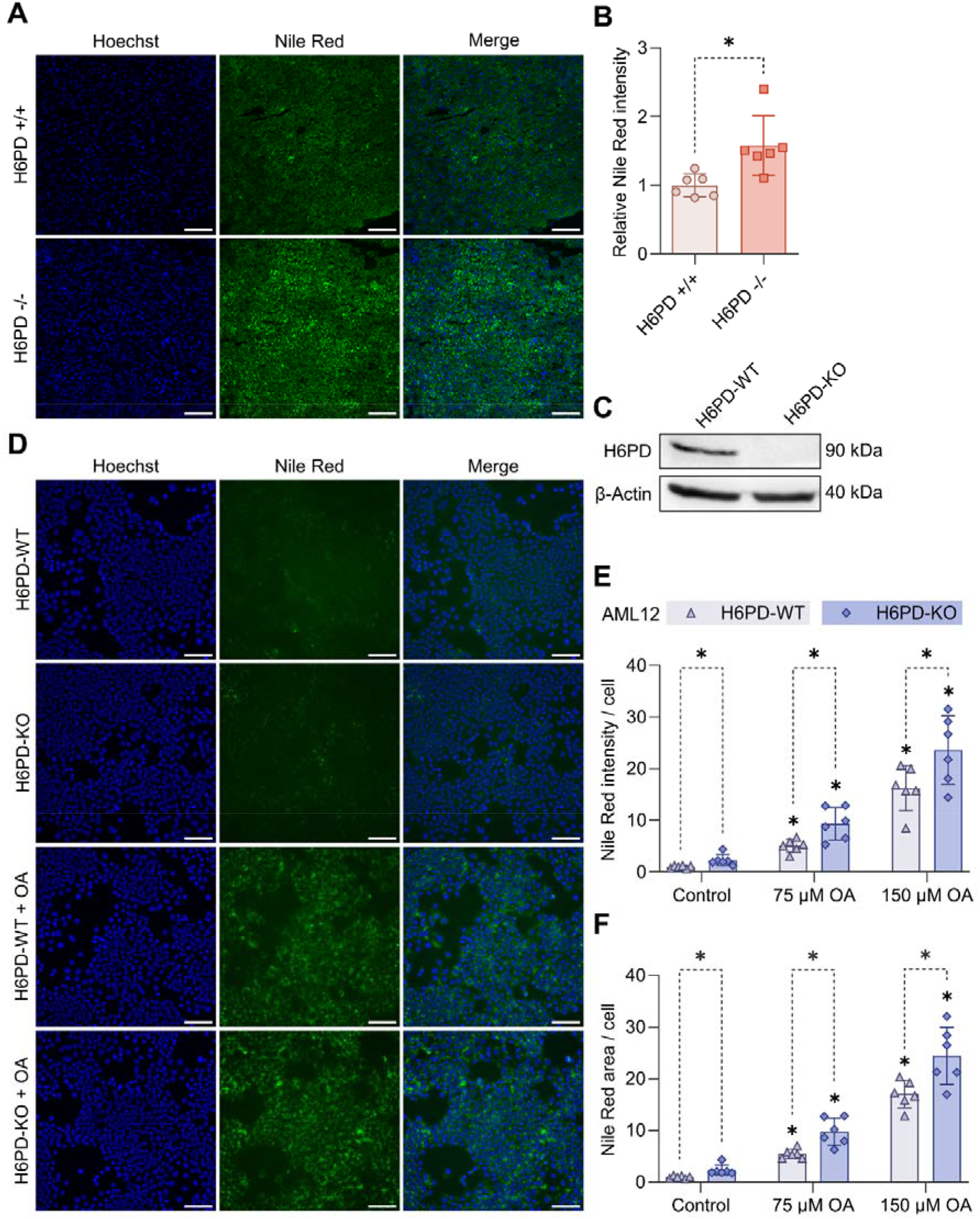
H6PD loss promotes neutral lipid accumulation in mouse liver tissue and AML12 hepatocytes. (**A**-**B**) Representative images of high-content imaging of Nile Red and Hoechst staining in H6PD+/+ and H6PD-/- mouse liver tissue sections (**A**) and corresponding quantification, normalized to the mean H6PD+/+ intensity (**B**). (**C**) Representative immunoblots of H6PD protein in AML12 H6PD-WT and -KO cells. β-actin served as loading control. (**D**) Representative images of high-content imaging of Nile Red and Hoechst staining in AML12 H6PD-WT and H6PD-KO cells cultured in standard medium with vehicle control (0.03% ethanol) or treated with 75 µM or 150 µM oleic acid (OA) for 48 h. (**E-F**) Quantification of Nile Red staining intensity (**E**) and lipid droplet area (**F**) normalized to the cell number. Scale bar = 100 μm. Data are presented as mean ± SD (n = 6 per group). Statistical significance was determined by two-way ANOVA followed by Fisher’s LSD post hoc test. *p < 0.05.

To confirm these observations *in vitro*, we generated an H6PD-KO AML12 murine hepatocyte cell line using CRISPR/Cas9-mediated genome editing. H6PD deletion was validated by Western blot analysis of AML12 cell lysates (Fig. 2C). Consistent with the *in vivo* findings, confocal microscopy of Nile Red- and Hoechst-stained cells revealed an accumulation of Nile Red-positive lipid droplets in H6PD-KO cells compared with H6PD-WT cells (Suppl. Fig. S3). This observation was confirmed by automated image acquisition and quantification demonstrating significantly increased Nile Red fluorescence intensity and lipid droplet area in H6PD-KO cells under basal conditions (Fig. 2D-F). Treatment with oleic acid at 75 µM or 150 µM for 48 h increased intracellular lipid accumulation in both genotypes; however, H6PD-KO cells consistently displayed greater Nile Red fluorescence intensity and lipid droplet area than H6PD-WT cells (Fig. 2E, F). Collectively, these findings demonstrate that H6PD deficiency promotes intracellular neutral lipid accumulation both in mouse liver and in AML12 cells, supporting a cell-autonomous role for H6PD in maintaining hepatic lipid homeostasis.

### 3.3. Proteomics-based pathway analysis reveals alterations in FA metabolism-associated proteins in H6PD-deficient liver

LC-HRMS-based proteomic analysis was conducted to further explore the consequences of H6PD loss in liver tissue and to generate hypotheses regarding the mechanisms underlying lipid accumulation in H6PD-/- mice (Fig. 3). In addition to liver homogenates, ER-enriched liver fractions (microsomes) were analyzed, aiming to improve the detection of ER-associated pathways. A total of 4569 and 3230 proteins were quantified in liver homogenates and microsomal fractions, respectively, from H6PD-/- and H6PD+/+ mice (Fig. 3A, 3B, Suppl. Table S4). As expected, H6PD was the most significantly downregulated protein in both fractions. GSEA of all differentially expressed proteins was conducted using the Hallmark gene set collection to identify biologically relevant patterns in the H6PD-/- proteome (Fig. 3C, 3D, Suppl. Table S4). Notably, fatty acid metabolism was among the most significantly enriched pathways in both liver homogenates and microsomal fractions and showed positive enrichment in H6PD-/- samples (Fig. 3C, 3D). In liver homogenates, peroxisomal pathways were highly upregulated as well, suggesting that the consequences of H6PD deficiency extend beyond the ER compartment (Fig. 3C). Additionally, the notable EMT signature may indicate alterations in cellular remodeling and tissue homeostasis as a response to the metabolic disruptions in the H6PD-/- mice. In microsomal fractions, the protein secretion gene set was negatively enriched, indicating potential disturbances in ER homeostasis. The enrichment of oxidative phosphorylation-related proteins may reflect a degree of mitochondrial contamination of the microsomal fraction and/or altered mitochondrial–ER interactions (Fig. 3D).

**Figure 3.**
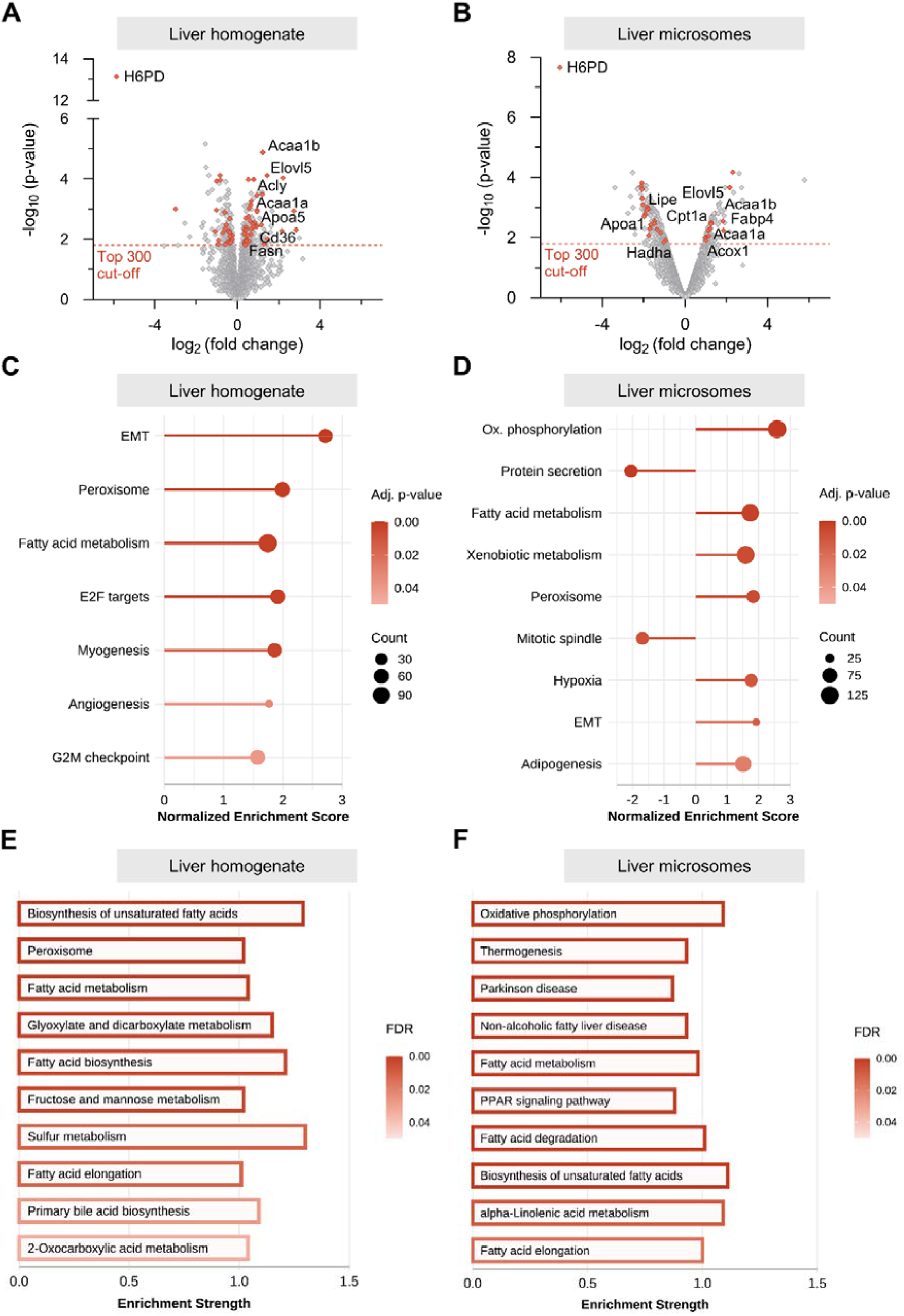
Proteomics-based pathway analysis of liver homogenates and microsomal fractions reveals dysregulation of lipid metabolism in H6PD-deficient mice. (A, **B)** Volcano plots representing the log2-fold change in protein abundance between H6PD-/- and H6PD+/+ liver homogenates (**A**) and liver microsomal fractions (**B**). The cut-off for the 300 most significantly altered proteins used for further analysis is indicated by red dotted line. Proteins associated with lipid metabolism are highlighted by red points, and exemplary proteins are designated by their names. (**C, D**) Significantly enriched pathways (adjusted p < 0.05) identified by GSEA of all differentially expressed proteins in liver homogenates (**C**) and microsomal fractions (**D**) using the Hallmark gene set collection. Pathways are ranked by adjusted P. EMT, epithelial-mesenchymal transition; ox. phosphorylation, oxidative phosphorylation. (**E, F**) The top 10 enriched pathways identified by KEGG pathway enrichment analysis of the 300 most significantly altered proteins using the STRING database in liver homogenates (**E**) and microsomal fractions (**F**). Pathways are ranked by FDR.

The 300 most significantly altered proteins (Fig. 3A, 3B) were further subjected to pathway analysis using the KEGG dataset via STRING (Fig. 3E, 3F, Suppl. Table S4). In liver homogenates, numerous lipid metabolism-related pathways, such as the biosynthesis of unsaturated fatty acids and fatty acid elongation, were among the most strongly enriched terms (Fig. 3E). While oxidative phosphorylation was again highly enriched in microsomes, pathways related to fatty acid synthesis, elongation, and degradation were also identified in this fraction (Fig. 3F).

Overall, H6PD loss led to a deregulation of proteins associated with multiple aspects of fatty acid metabolism (Fig. 3A, Suppl. Table S4). More specifically, this included a significant upregulation of several enzymes involved in peroxisomal fatty acid β-oxidation, such as ATP-binding cassette subfamily D member 2 (ABCD2, log2FC = 2.740 for liver homogenate) and 3-ketoacyl-CoA thiolase B (ACAA1B, log2FC = 1.229) as well as peroxisomal and mitochondrial acyl-CoA thioesterase family members (ACOT3 log2FC = 2.206; ACOT2, log2FC = 0.955). Additionally, proteins involved in fatty acid uptake, including cluster of differentiation 36 (CD36, log2FC = 0.924) and, to a lesser degree, fatty acid-binding protein 1 (FABP1, log2FC = 0.572) were increased in H6PD-/- liver homogenates. Interestingly, ER-associated enzymes involved in fatty acid elongation and desaturation, such as elongation of very long-chain fatty acids protein 5 (ELOVL5, log2FC = 1.431) and stearoyl-CoA desaturase 1 (SCD1, log2FC = 1.203), were also upregulated in KO mice. Together, these alterations indicate a broad metabolic adaptation involving enhanced FA uptake, processing, and remodeling pathways in response to H6PD deficiency. However, despite the induction of these compensatory pathways, lipid accumulation persists in H6PD-/- livers, suggesting that these responses are insufficient to restore hepatic lipid homeostasis.

### 3.4. H6PD-KO AML12 cells preferentially rely on FAs as mitochondrial fuel but exhibit increased susceptibility to lipotoxicity

Building on the proteomic findings, a series of functional assays were conducted to compare lipid handling between H6PD-WT and -KO AML12 cells. First, mitochondrial fuel dependency on the main metabolic substrates glutamine, glucose, and FAs was evaluated using a Seahorse Mito Fuel Flex assay (Fig. 4A). H6PD-KO cells exhibited a marked increase in dependency on FA oxidation, with FAs representing the predominant mitochondrial fuel source. In contrast, H6PD-WT cells showed greater dependency on glucose, whereas glucose was the least utilized substrate in H6PD-KO cells. Glutamine dependency was also increased in H6PD-KO cells compared to H6PD-WT cells (Fig. 4A). Given the altered substrate utilization observed in H6PD-KO cells, we next investigated whether H6PD deficiency also affects glycolytic function. A Glycolysis Stress Test was performed, in which glucose, oligomycin, and 2-deoxyglucose (2-DG) were sequentially injected to determine glycolytic rate, glycolytic capacity, and glycolytic reserve (Fig. 4B). No significant differences were observed between H6PD-WT and H6PD-KO cells in any of these parameters, indicating that H6PD deficiency does not substantially alter glycolytic capacity under the conditions tested.

**Figure 4.**
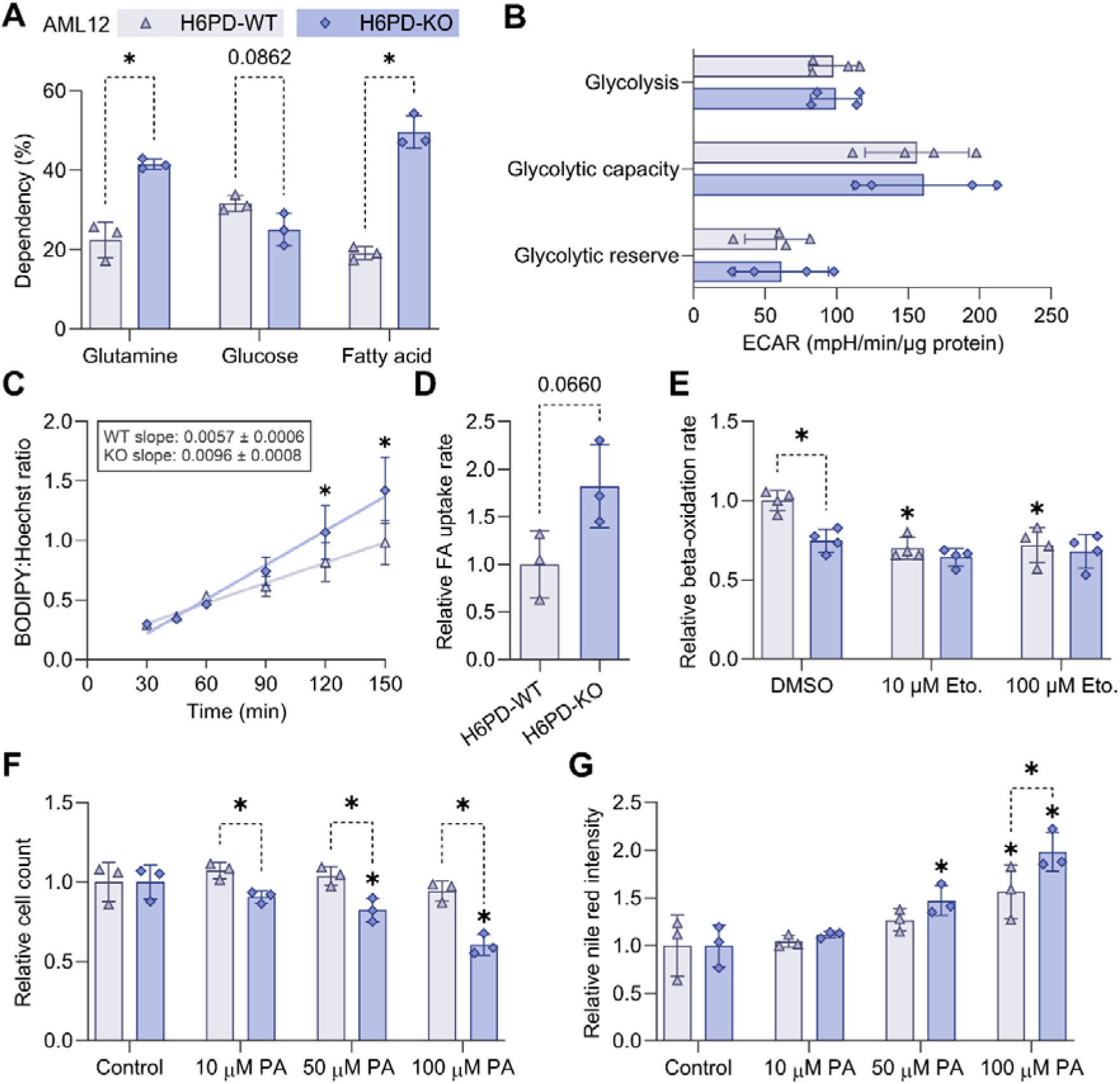
H6PD-deficient AML12 cells exhibit increased dependency on FAs as mitochondrial fuel but increased susceptibility to PA-induced lipotoxicity. **(A)** Mitochondrial substrate dependency on glucose, glutamine, and FAs was assessed in H6PD-WT and H6PD-KO AML12 cells using the Mito Fuel Flex Test. **(B)** Glycolytic parameters, including glycolytic capacity and glycolytic reserve, were calculated from the Glycolysis Stress Test data. **(C)** Relative FA uptake in AML12 cells after incubation with BODIPY-C16. Mean fluorescence intensity at the indicated time points after treatment, normalized to Hoechst-33258 intensity. Linear regression slopes and corresponding p values are indicated. **(D)** Normalized slopes from each biological replicate were determined to compare FA uptake rates. **(E)** FA β-oxidation rates were determined by measuring the oxidation of radiolabeled palmitate. The specificity of this assay was confirmed using etomoxir (10 and 100 µM). (**F**) Cell number after 48 h of treatment with palmitic acid (PA) normalized to the corresponding vehicle control (0.08% ethanol). (**G**) Intracellular neutral lipid accumulation following PA treatment was assessed by Nile Red staining and high-content imaging. Nile Red fluorescence intensity was normalized to cell number and the corresponding vehicle control. Data are presented as mean ± SD (n = 3). Statistical significance was determined by Welch’s t test or two-way ANOVA followed by Fisher’s LSD post hoc test. *p < 0.05.

Next, FA uptake was assessed by incubating cells with fluorescently labeled palmitic acid and quantifying the intracellular fluorescence intensity following fixation at increasing time points (Fig. 4C). Comparison of the linear regression slopes revealed a trend toward higher FA uptake rates in H6PD-KO cells compared with H6PD-WT (Fig. 4D). As the substrate dependency test suggested an increased reliance on FAs in H6PD-KO cells, we further investigated mitochondrial fatty acid β-oxidation in both cell models. Cells were incubated with radiolabeled palmitate for 3 h to measure palmitate oxidation, using the carnitine palmitoyltransferase I (CPT1) inhibitor etomoxir as a pharmacological control (Fig. 4E). As expected, etomoxir at 10 and 100 µM effectively inhibited palmitate oxidation in H6PD-WT cells, confirming the experimental setup. Residual β-oxidation activity may reflect the contribution of peroxisomal FA oxidation pathways. Remarkably, under basal conditions, the rate of mitochondrial β-oxidation was significantly lower in H6PD-KO cells compared to H6PD-WT cells.

Finally, susceptibility to lipotoxicity was assessed by exposing cells to BSA-coupled PA at concentrations ranging from 10 to 100 µM (Fig. 4F, 4G). Cell number was quantified following Hoechst staining using a Cytation 5 imaging reader. Notably, at all PA concentrations tested, H6PD-KO cells displayed significantly lower cell numbers than H6PD-WT cells (Fig. 4F). This difference was most marked at the highest PA concentration tested. In parallel, intracellular TG accumulation in response to the PA treatment was evaluated by Nile Red staining (Fig. 4G). H6PD-KO cells showed a trend toward a greater relative increase in Nile Red fluorescence following PA treatment, reaching statistical significance at the highest PA concentration tested.

Collectively, these findings indicate that H6PD deficiency induces a metabolic shift toward enhanced FA utilization. Despite this apparent adaptation, H6PD-KO cells displayed increased sensitivity to PA-induced loss of cell number, suggesting that increased FA uptake and altered FA utilization may be accompanied by an impaired capacity to cope with excess lipid load.

### 3.5. H6PD deficiency is associated with altered CPT1 expression

Given the altered FAO rate in H6PD deficiency models, we further investigated the functional integrity of this pathway. The transport of long-chain FAs into the mitochondrial matrix as acyl-CoA derivatives depends on the carnitine shuttle system, mediated by CPT1 and CPT2, located at the outer mitochondrial membrane and the matrix side of the inner mitochondrial membrane, respectively (45). We first measured their protein abundance by immunoblot analysis. In H6PD-KO cells, CPT1A and CPT1B protein levels were significantly reduced compared with their respective controls, while CPT2 levels showed a modest decrease that did not reach statistical significance (Fig. 5A, 5B). A similar reduction in CPT1 levels was observed in livers from H6PD-/- mice (Suppl. Fig. S4A, B).

**Figure 5.**
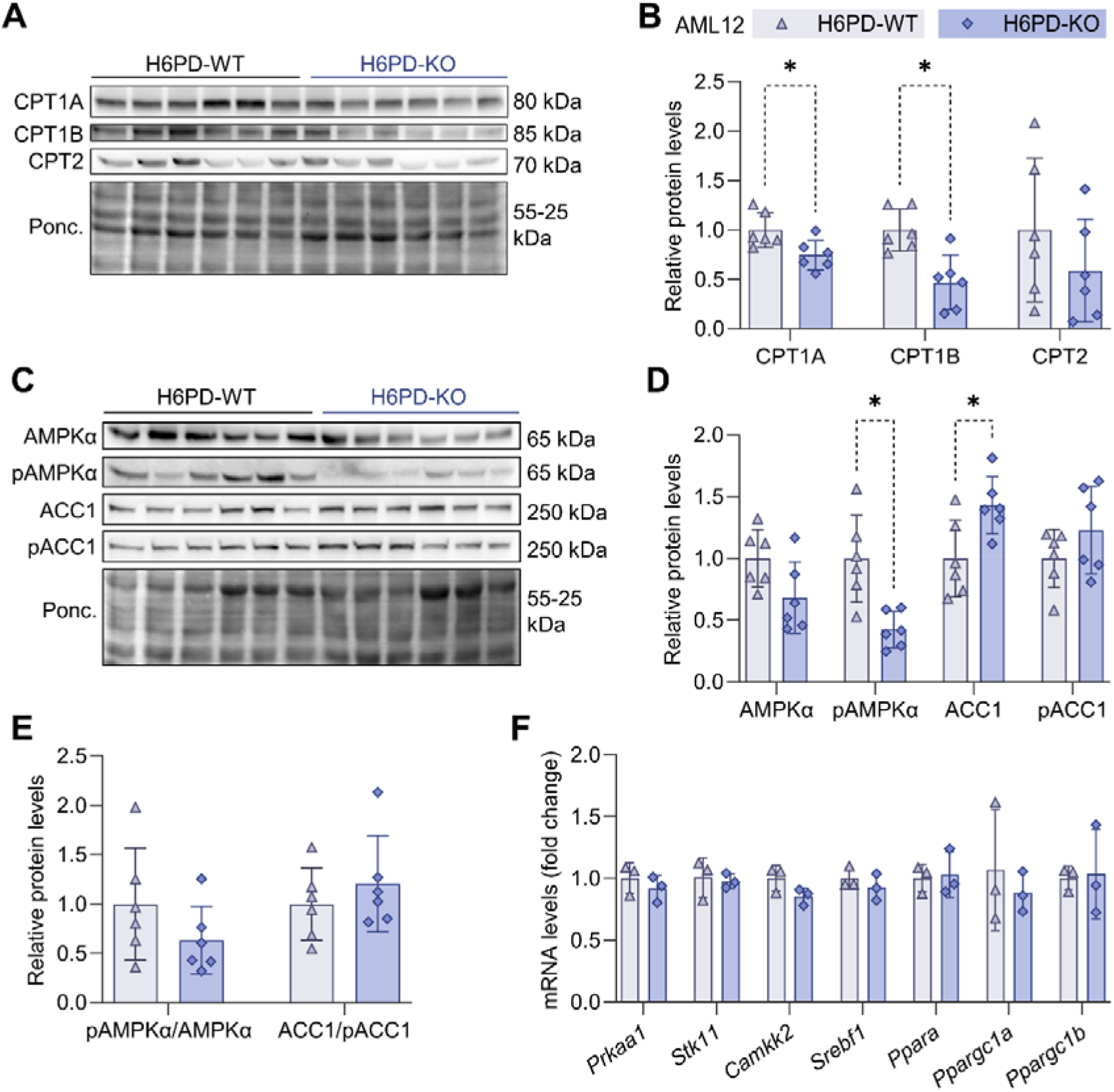
CPT1 protein levels are decreased in H6PD-deficient mice livers and AML12 cells, concomitant with reduced mtFAO. (**A**) Immunoblots of CPT1A, CPT1B, and CPT2 and representative Ponceau staining of AML12 H6PD-WT and -KO cells. (**B**) Protein levels, calculated by densitometry and normalized to Ponceau staining, are displayed relative to the respective control group. (**C**) Immunoblots of AMPKα, pAMPKα (Thr172), ACC1, and pACC1 (Ser79). (**D, E**) Protein levels, calculated by densitometry and normalized to Ponceau staining, are displayed relative to the respective control group. (**F**) mRNA levels normalized to *18S* and shown as fold change relative to their respective control group. Data are presented as mean ± SD (n = 6 for immunoblots, n = 3 for RT-qPCR). Statistical significance was determined by an unpaired two-tailed t-test with Welch’s correction. *p < 0.05.

Additionally, we determined protein levels and the activation status of acetyl-CoA carboxylase (ACC), which can inhibit CPT1 through the production of malonyl-CoA in cells and liver samples (Fig. 5C-E, Suppl. Fig. S4C-E, respectively). Total levels of ACC1, the isoform predominantly expressed in liver, were modestly increased in H6PD-KO cells. Levels of phosphorylated (Ser79) and therefore inactivated, ACC1, as well as the ACC1/pACC1 ratio, were not significantly altered (Fig. 5C-E). In liver samples, no alteration in the ACC1/pACC1 ratio was detected either (Suppl. Fig. S4C-E).

Since ACC is regulated by 5′-AMP-activated protein kinase (AMPK), a major cellular energy sensor and metabolic regulator (46), we additionally measured protein levels and activation status of its catalytic subunit AMPKα (Fig. 5C-E). We observed a reduction of total AMPKα protein in H6PD-KO cells that did not reach significance and a significant reduction in pAMPKα levels. The pAMPKα/AMPKα ratio was not significantly altered. Although AMPKα levels were significantly reduced in mouse liver, there was no difference in AMPKα activation status in the liver tissue either (Suppl. Fig. S4C-E).

To further explore mechanisms involved in coordinating cellular responses to H6PD deficiency, we performed RT-qPCR experiments using the AML12 cell lines (Fig. 5F). We detected no significant decrease in the mRNA levels of *Prkaa1* (encoding for AMPKα1) or its regulators *Stk11* and *Camkk2* in H6PD-KO. There was also no alteration in the gene expression of other key regulators of hepatic metabolism, namely *Srebf1* and *Ppara*. We observed greater variability in the gene expression of *Ppargc1a* and *Ppargc1b,* but no significant difference between the H6PD-WT and H6PD-KO lines (Fig. 5F). Due to the high biological variability in RNA isolated from mouse tissue, we could not reliably perform these experiments with liver samples.

Collectively, these findings identify an impairment of hepatic lipid metabolism associated with H6PD deficiency, which may contribute to the observed accumulation of hepatic TGs. Although there may be some disturbances in the AMPK/ACC axis in models of H6PD deficiency, these changes are unlikely to fully explain the observed metabolic alterations, suggesting that alternative pathways are involved in sensing the loss of H6PD and orchestrating the resulting hepatic metabolic rewiring.

## 4. Discussion

In the present study, we integrated proteomic, lipidomic, and functional analyses to investigate the consequences of H6PD deficiency in mouse liver tissue and AML12 hepatocytes. Our findings demonstrate that the loss of H6PD leads to profound remodeling of hepatic lipid metabolism, characterized by the selective accumulation of long-chain, highly unsaturated TG species and overall intracellular neutral lipid accumulation. Proteomic analysis revealed increased expression of proteins involved in hepatic FA handling in H6PD-deficient mice, while functional assays demonstrated impaired mtFAO in H6PD-KO AML12 hepatocytes. Together, these results identify H6PD as an important regulator of hepatic lipid homeostasis and suggest that H6PD-dependent maintenance of ER luminal redox homeostasis may be important for efficient mitochondrial lipid utilization.

A major finding of our lipidomic analysis was the structural selectivity of the altered TG pool in H6PD-/- livers. Rather than a global, uniform elevation of all lipid classes, H6PD ablation selectively promoted the accumulation of long-chain, highly polyunsaturated TG species alongside oxTGs. Concurrently, we observed marked reductions in several phospholipid classes, specifically PC, LPE, and LPC. The depletion of PC, the major phospholipid component of the lipid droplet monolayer (47), together with reduced LPC levels, suggests altered phospholipid remodeling that may influence lipid droplet dynamics and hepatic lipid export pathways, including VLDL assembly and secretion (48). Notably, increased accumulation of long-chain and unsaturated fatty acids in H6PD-deficient livers was accompanied by enhanced lipid droplet formation, consistent with the concept that hepatocytes mitigate excess free FA toxicity by esterifying FA into TGs for storage within lipid droplets, which originate from the ER membrane (49).

The structural shift toward long-chain, unsaturated TG species is consistent with the increased expression of the ER-associated elongase ELOVL5 and the desaturase SCD1, suggesting activation of adaptive lipid remodeling pathways. While these enzymatic adaptations may protect hepatocytes by channeling potentially lipotoxic FA into neutral TG stores (50), the enrichment of highly polyunsaturated acyl chains may increase the susceptibility of the hepatic lipid pool to lipid peroxidation, consistent with the elevated levels of oxTGs observed in H6PD-deficient mouse livers.

Our functional characterization revealed a distinct metabolic paradox in H6PD-deficient hepatocytes. Substrate dependency assays demonstrated that H6PD-KO cells showed increased mitochondrial dependency on FA oxidation relative to glucose. However, direct flux assays revealed that their mitochondrial β-oxidation capacity was significantly diminished. This apparent contradiction is reconciled by the fact that the two assays probe different aspects of FA metabolism. The substrate-dependency assay reflects how much cells currently draw on endogenous intracellular FA stores under baseline conditions, whereas the direct β-oxidation assay is performed in the presence of exogenous FA and therefore captures oxidative capacity under substrate-saturating, near-maximal conditions. Thus, the data indicate that the increased FA dependency in H6PD-KO cells does not necessarily indicate enhanced β-oxidation capacity, but rather a greater reliance on FA-derived substrates despite an impaired capacity to oxidize them. This discrepancy between substrate dependency and oxidative capacity creates a metabolic bottleneck: hepatocytes increase FA uptake (supported by the proteomic upregulation of CD36) and rely more heavily on FA fuel pathways, yet exhibit a reduced capacity for mtFAO. This imbalance may promote the redirection of excess FAs toward TG synthesis and intracellular storage, which may contribute to the increased Nile Red-positive lipid droplet area and the heightened susceptibility of H6PD-deficient cells to palmitic acid-induced lipotoxicity.

At the molecular level, the impaired mtFAO observed in H6PD-deficient hepatocytes was associated with a reduction in CPT1 protein expression, a key component of the mitochondrial import pathway of long-chain fatty acyl-CoAs. Interestingly, this decrease in CPT1 abundance is unlikely to be explained solely by alterations in the canonical AMPKα/ACC signaling pathway. Although total AMPKα protein levels were moderately reduced, phosphorylation of AMPKα (Thr172) and its downstream target ACC (Ser79), normalized to total protein abundance, remained largely unchanged in both H6PD-deficient AML12 cells and mouse livers. The transcriptional regulation of CPT1 is also controlled by mechanisms independent of the AMPKα/ACC axis. In particular, PPARα and its transcriptional coactivator PGC-1α are major regulators of genes involved in mtFAO, including CPT1 (51). Although we did not detect changes in the mRNA expression of either *Ppara* or *Ppargc1a*, the activity of PGC-1α can be regulated through post-translational mechanisms, including phosphorylation by AMPK and other signaling pathways, rather than by changes in gene expression alone (52). Consequently, altered PGC-1α activity cannot be excluded as a contributor to the reduced expression of CPT1 in H6PD-deficient hepatocytes. Alternatively, the reduction in CPT1 expression may represent a secondary adaptation to chronic lipid accumulation and metabolic remodeling induced by H6PD deficiency. In this context, hepatocytes may limit mitochondrial FA import while redirecting excess fatty acids toward TG synthesis, lipid droplet storage, and peroxisomal β-oxidation, thereby reducing the burden on an already compromised mitochondrial oxidative system. This interpretation is not only consistent with the observed TG accumulation and enhanced lipid droplet formation, but also the broad induction of peroxisomal fatty acid oxidation pathways in the proteomics analysis of H6PD-deficient livers. Key peroxisomal proteins involved in fatty acid transport and oxidation, including ABCD2, ACAA1B, ACOT2, and ACOT3, were increased in H6PD-deficient livers, suggesting that enhanced peroxisomal processing may represent an adaptive response to compensate for reduced mitochondrial fatty acid utilization. Peroxisomes are particularly important for the initial shortening of VLCFAs, which are subsequently transferred to mitochondria for complete oxidation (8). Thus, H6PD-deficient hepatocytes may increase their reliance on peroxisomal fatty acid oxidation to facilitate the disposal of excess long-chain and VLCFA accumulated in the liver. However, this compensatory response may be incomplete, as peroxisomal β-oxidation does not directly produce ATP and generates hydrogen peroxide as a byproduct. Consequently, increased reliance on peroxisomal oxidation may reduce lipid overload while potentially increasing oxidative stress and failing to fully restore mitochondrial energy metabolism. Together, these findings highlight the metabolic flexibility of H6PD-deficient hepatocytes but also suggest that activation of alternative lipid disposal pathways may contribute to redox imbalance and hepatic lipid dysfunction.

Because H6PD is localized exclusively within the ER lumen, the mitochondrial defects observed in our models highlight a potential role for inter-organellar communication. In our microsomal proteomic analyses, pathways related to oxidative phosphorylation were notably enriched. Although some mitochondrial protein detection may reflect unavoidable microsomal contamination, enrichment of oxidative phosphorylation pathways is also consistent with altered ER–mitochondrial interactions or remodeling of mitochondria-associated ER membranes (MAMs), which are known to coordinate metabolic cross-talk, lipid transfer, and bioenergetics under ER stress conditions (53). A recent study has linked H6PD to MAM integrity, mitochondrial redox homeostasis and mitophagy in HeLa human cervical cancer cells (54). Elucidating how disruption of ER luminal NADPH homeostasis is transmitted to mitochondria therefore represents an important direction for future research.

Presently, H6PD is the only known enzyme that generates NADPH within the lumen of the ER. Although H6PD is indispensable for 11β-HSD1 reductase activity (29), liver-specific deletion of 11β-HSD1 does not reproduce the hepatic TG accumulation or major alterations in lipid metabolism observed in our models (55). Moreover, studies in breast cancer cells and skeletal muscle have shown that H6PD deficiency induces marked metabolic alterations independently of 11β-HSD1 (56, 57), suggesting that additional H6PD-dependent pathways remain to be identified. Likewise, our recent BioID-based interactome study localized H6PD within a protein network associated with oxidative protein folding and ER proteostasis, although no direct NADPH-dependent protein targets were identified (58). These observations support the notion that H6PD contributes to maintaining ER homeostasis beyond glucocorticoid metabolism. An attractive candidate linking H6PD activity to hepatic lipid metabolism is the microsomal enzyme 17β-hydroxysteroid dehydrogenase 12 (17β-HSD12/KAR) (59), which catalyzes the reduction step during elongation of long-chain fatty acids (19). Its predicted luminal orientation and preference for NADPH make it a plausible candidate for consuming H6PD-derived reducing equivalents. Consistent with this possibility, hepatocyte-specific deletion of 17β-HSD12 results in hepatic steatosis and liver injury, phenotypes that partially resemble those observed in our H6PD-deficient models (60). Furthermore, both studies identified marked reductions in major urinary proteins, particularly MUP1, a hepatokine whose expression is suppressed during ER stress and metabolic dysfunction (61). Although our proteomic data did not reveal altered expression of 17β-HSD family members, these phenotypic similarities suggest that disruption of ER luminal NADPH availability may influence lipid homeostasis through pathways involving, but not necessarily limited to, 17β-HSD12. Whether 17β-HSD12 directly depends on H6PD-derived NADPH remains an important question for future studies.

### 4.1. Conclusions

Our study identifies H6PD as an important regulator of hepatic lipid homeostasis and demonstrates the impaired mtFAO as a central functional consequence of H6PD deficiency. Rather than acting solely through the canonical 11β-HSD1 pathway or through canonical AMPK-ACC signaling, loss of H6PD triggers a broad metabolic rewiring that links H6PD deficiency and disruption of ER luminal redox homeostasis with mitochondrial oxidative dysfunction and polyunsaturated TG accumulation. These findings reveal an unexpected connection between ER NADPH homeostasis and hepatic energy metabolism, providing a conceptual framework for identifying the molecular signals that coordinate metabolic communication between the ER and mitochondria.

### 4.2. Limitations

One of the main limitations of this study is the exclusive use of male mice. Previous studies have demonstrated sex-specific differences in hepatic lipid metabolism and susceptibility to steatotic liver disease that may limit the applicability of our findings across sexes (62, 63). Future studies should include both sexes to determine whether H6PD deficiency has sex-dependent effects.

Another limitation is that all animals were maintained on a standard chow diet. Challenging H6PD-/- mice with metabolic stressors, such as high-fat or Western diets, would help determine whether H6PD loss exacerbates steatohepatitis and liver injury. In addition, the use of a global H6PD KO model does not entirely eliminate the potential influence of extrahepatic systemic signals on the liver phenotype, although our *in vitro* findings in AML12 cells support a cell-autonomous role for H6PD in hepatocytes. Finally, translational validation using human hepatocyte models is required to assess the relevance of the H6PD to human hepatic lipid metabolism and metabolic disease.

## Data availability

The data generated and/or analyzed in the current study will be made available in the Zenodo repository.

## Supplemental data

This article contains supplementary data.

## Acknowledgements

We thank Diego Calabrese (Department of Biomedicine, Histology Core Facility, University of Basel, Switzerland) for his support in histology.

## Author contributions

**Gabriele Sakalauskaite**: Conceptualization, Formal analysis, Investigation, Data Curation; Writing – Original Draft, Visualization; **Sophie Ebert**: Formal analysis, Investigation, Data Curation; Writing – Original Draft, Visualization; **Julien Arthur Allard**: Conceptualization, Methodology; **Michael Zogg**: Formal analysis, Investigation; **Isabel Meister**: Formal analysis, Investigation, Data Curation; Visualization; **Julia Birk**: Methodology; **Alexander Schmidt**: Supervision, Data Curation; Formal analysis; **Jamal Bouitbir**: Conceptualization, Supervision, Writing – Original draft, Visualization; **Alex Odermatt**: Conceptualization, Supervision, Funding acquisition; Project Administration; Writing – Original Draft, Visualization.

## Abbreviations

ABCD2: ATP-binding cassette subfamily D member 2
ACC: acetyl-CoA carboxylase
ACAA1B: 3-ketoacyl-CoA thiolase B
ACOT: acyl-CoA thioesterase
AMPK: 5′-AMP-activated protein kinase
CPT1/2: carnitine palmitoyltransferase I/II
CD36: cluster of differentiation 36
ECAR: extracellular acidification rate
ELOVL5: elongation of very long-chain fatty acids protein 5
EMT: epithelial-mesenchymal transition
FABP1: fatty acid-binding protein 1
FDR: false discovery rate
G6P: glucose-6-phosphate
GSEA: gene set enrichment analysis
H6PD: hexose-6-phosphate dehydrogenase
KAR: 3-ketoacyl-CoA reductase
LC-HRMS: liquid chromatography–high-resolution mass spectrometry
LPC: lysophosphatidylcholine
LPE: lysophosphatidylethanolamine
LSD: least significant difference
MAM: mitochondria-associated endoplasmic reticulum membrane
MASLD: metabolic dysfunction-associated steatotic liver disease
mtFAO: mitochondrial fatty acid oxidation
oxTG: oxidized triacylglycerols
PC: phosphatidylcholine
PFA: paraformaldehyde
PPP: pentose phosphate pathway
SCD1: stearoyl-CoA desaturase 1
TECR: trans-2-enoyl-CoA reductase
TG: triglyceride
UPR: unfolded protein response
VLCFA: very-long-chain fatty acid
11β-HSD1: 11β-hydroxysteroid dehydrogenase type 1
2-DG: 2-deoxyglucose.

