## Supplemental Figure S1, S2, S3, and S4; Supplemental Table S1, S2, and S3. for "Hexose-6-phosphate dehydrogenase deficiency disrupts hepatic fatty acid homeostasis and induces triglyceride accumulation"

**Supplemental data**

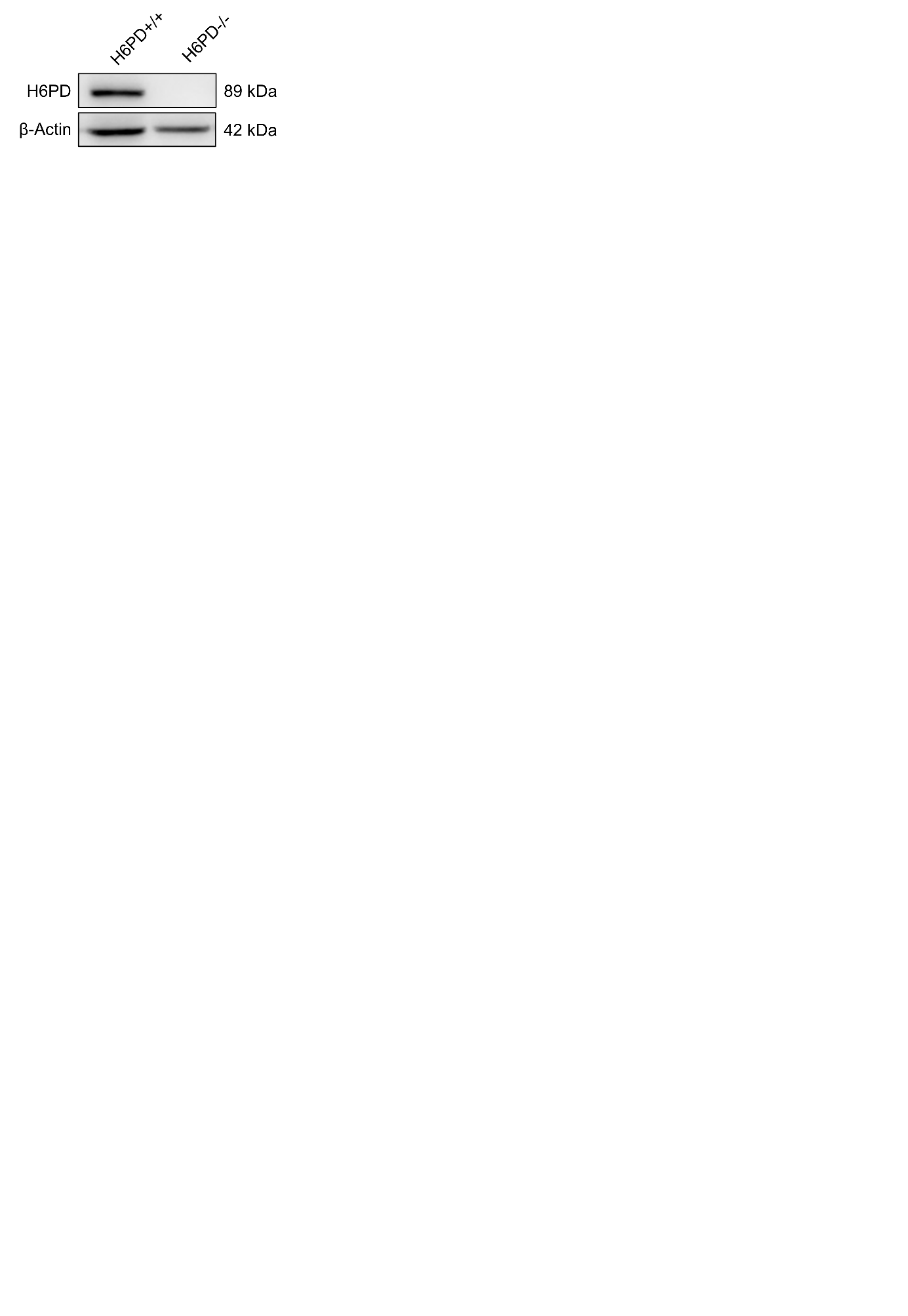

**Supplemental Figure S1. Confirmation of H6PD KO in the H6PD-/-** **C57BL/6J mice.** Representative Western blot showing H6PD levels in liver tissue from H6PD+/+ and H6PD-/- mouse. β-Actin served as a loading control.

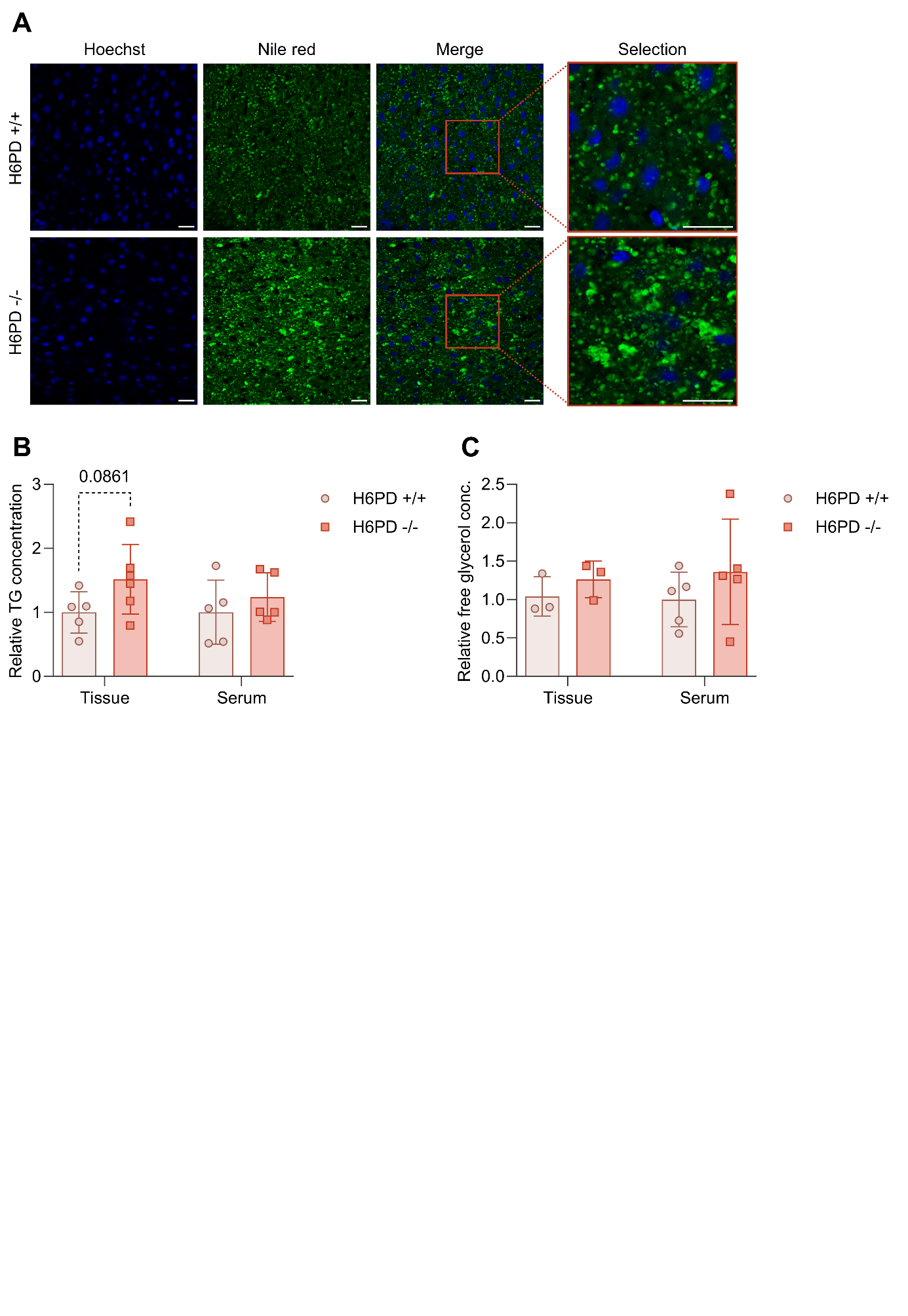

**Supplemental Figure S2. Nile Red staining and TG measurement of mouse liver tissue and serum.** (**A**) Representative images of Nile Red and Hoechst stainings in H6PD+/+ and H6PD-/- mouse liver tissue, acquired by confocal microscopy. Scale bar = 20 μm. (**B**) Relative triglyceride and (**C**) relative free glycerol concentrations in H6PD+/+ and H6PD-/- mouse liver tissue and serum. Measurements are normalized to sample protein concentration for liver tissue or sample volume for serum, and to the H6PD+/+ mean. Data are presented as mean ± SD (n = 6 except for free glycerol measurements in liver tissue, where n = 3). Statistical significance was determined by unpaired two-tailed t-test with Welch’s correction. *p < 0.05.

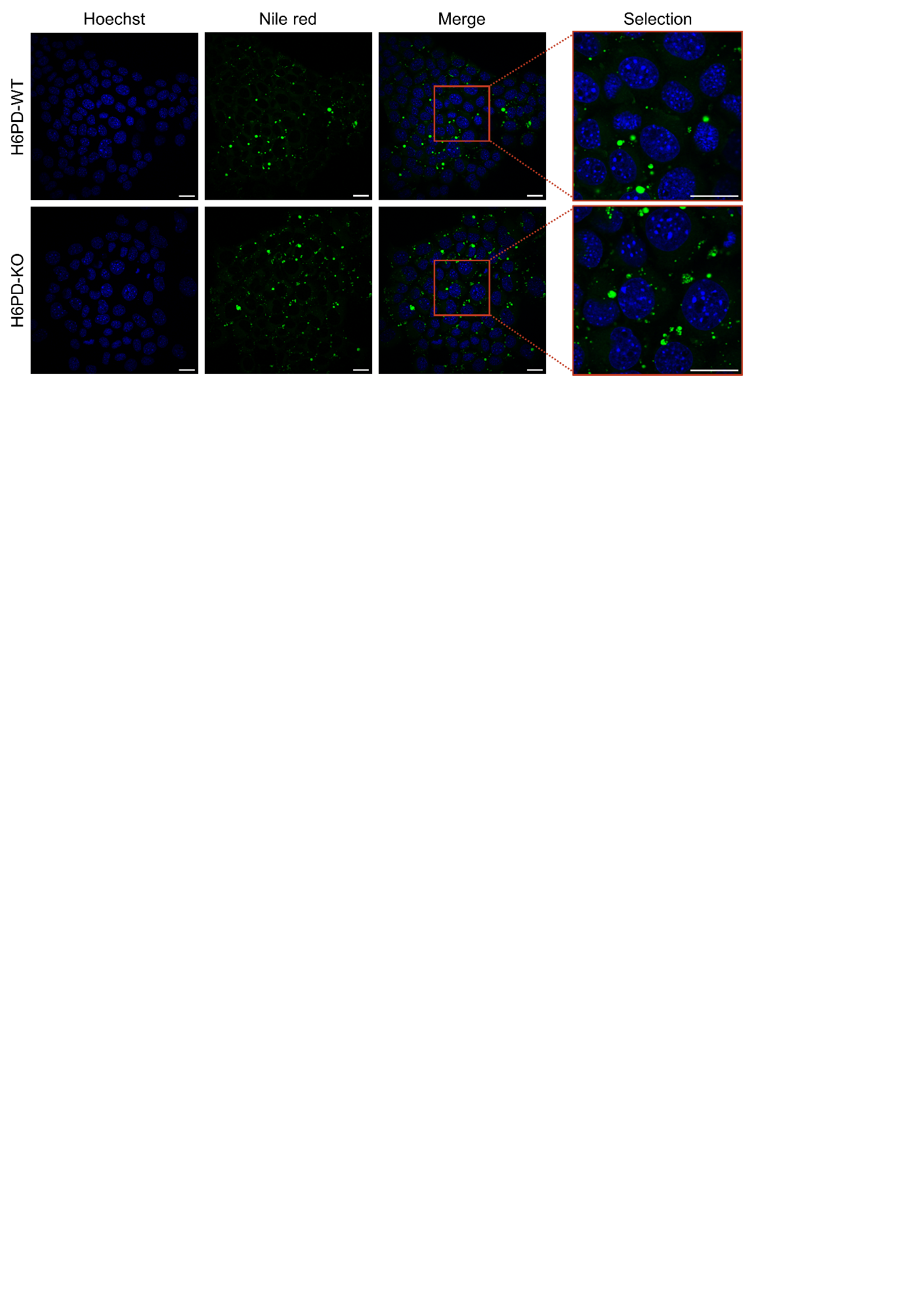

**Supplemental Figure S3. Nile Red staining of AML12 cells.** Representative images of Nile Red and Hoechst staining in AML12 H6PD-WT and H6PD-KO cells, acquired by confocal microscopy. Scale bar = 20 μm.

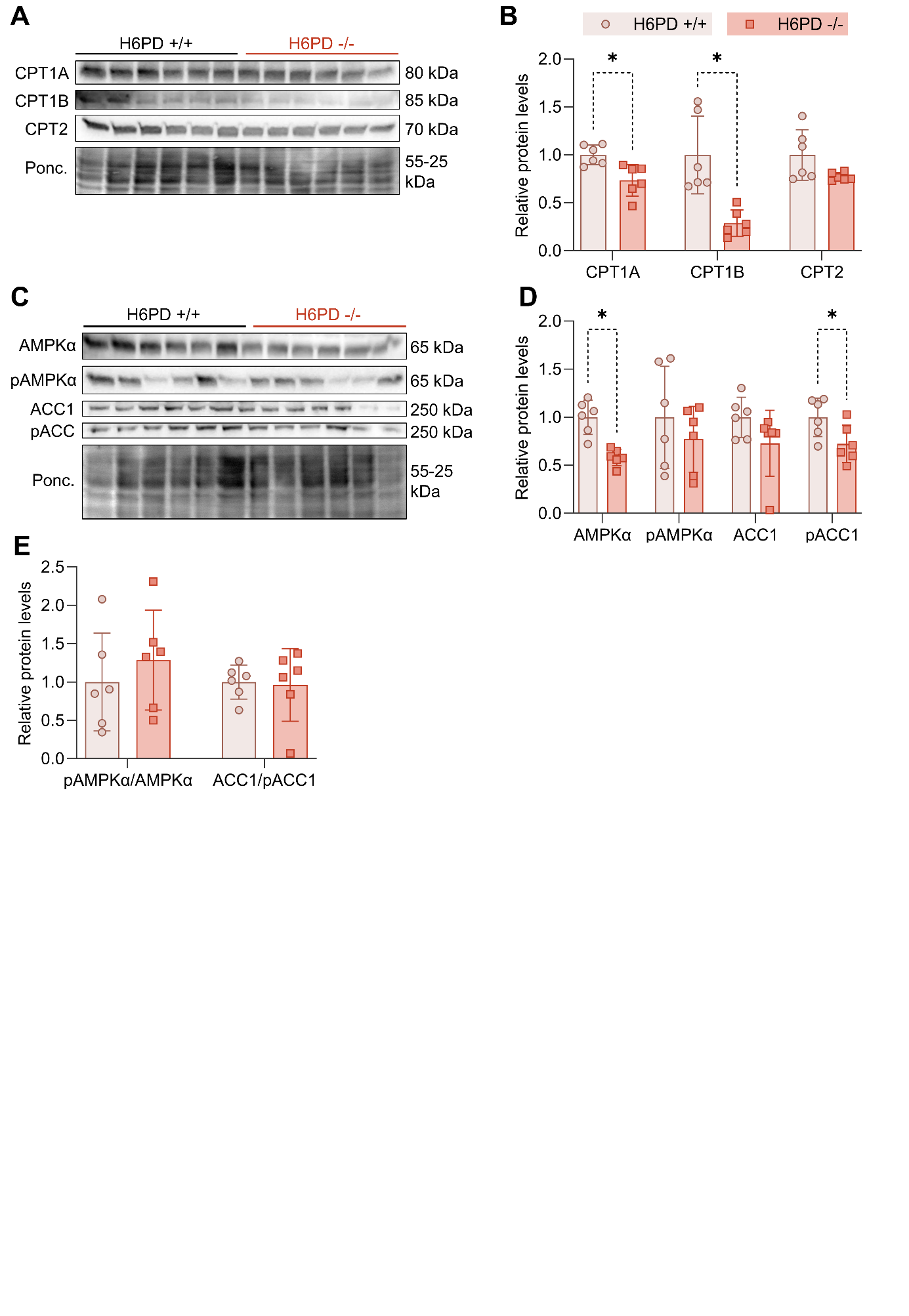

**Supplemental Figure S4. CPT1 and AMPK**α **protein levels in mouse liver tissue.** (**A**) Immunoblots of CPT1A, CPT1B, and CPT2 and representative Ponceau staining of H6PD+/+ and H6PD-/- mouse liver tissue. (**B**) Protein levels calculated by densitometry and normalized to Ponceau staining are displayed relative to the respective control group. (**C**) Immunoblots of AMPKα, pAMPKα (Thr172), ACC1, and pACC1 (Ser79). (**D, E**) Protein levels calculated by densitometry and normalized to Ponceau staining are displayed relative to the respective control group. Statistical significance was determined by an unpaired two-tailed t-test with Welch’s correction. *p < 0.05.

| Name | Abbreviation | Catalog number | Company |
| --- | --- | --- | --- |
| **DL-Dithiothreitol** | DTT | **#D9163** | Sigma-Aldrich, Darmstadt, Germany |
| **Tris(2-carboxyethyl)phosphine hydrochloride** | TCEP | **#C4706** |  |
| **Oleic acid** | OA | **#O100-8** |  |
| **Trypan blue** |  | **#T8154** |  |
| **Methanol** |  | **#32213** |  |
| **Trizma® base** |  | **#1003471714 T1503** |  |
| **Glycine** |  | **#G-7126** |  |
| **Ammoniumpersulfate** | APS | **#A3678** |  |
| **Tetramethylenediamine** | **TEMED** | **#T9281** |  |
| **Methanol** |  | **#32213** |  |
| **Triethyloammonium bicarbonate** | **TEAB** | **#18597** |  |
| **Phosphoric acid** |  | **#438081** |  |
| **Sodium dodecyl sulfate** | **SDS** | **#71725** |  |
| **Iodoacetamide** |  | **#I1149** |  |
| **Formic acid** |  | **#94318** |  |
| **Nile Red** |  | **#N3013** |  |
| **Urea** |  | **#1.08487.1000** | Merck, Darmstadt, Germany |
| **Palmitic acid** |  | **#P0500** |  |
| **Paraformaldehyde** | **PFA** | **#4005** |  |
| **Perchloric acid, 60%** |  | **#**311413 |  |
| **Phosphatase inhibitor cocktail PhosSTOP** |  | **#04906837001** | **F. Hoffmann-La Roche, Basel, Switzerland** |
| **cOmplete Mini Protease Inhibitor cocktail tablets** | **PIC** | **#11836153001** |  |
| **Bovine Serum Albumin Fraction V** | **BSA** | **#10735108001** |  |
| **Acrylamide** |  | **#A1672,1000** | **PanReac AppliChem,** Darmstadt Germany |
| **Sodium Chloride** |  | **#A2942,5000** |  |
| **Sodium dodecyl sulfate** | **SDS** | **#9T0122920** |  |
| **Porcine Trypsin** |  | **#V5113** | **Promega, Fitchburg, WI, USA** |
| **HPLC water** |  | **#W/0106/17** | **Chemie Brunswick, Basel, Switzerland** |
| **Acetonitrile** |  | **#A955-212** | **Thermo Scientific, Waltham, MA, USA** |
| **Tween® 20** |  | **#0777-1L** | **VWR Chemicals Solon Ohio, Solon, OH, USA** |
| **Milk Powder** |  | **#1929** | **COOP, Bern, Switzerland** |
| **Radioimmunoprecipitation assay Buffer** | **RIPA** | **#39244.02** | **SERVA, Heidelberg, Germany** |

**Supplemental Table S1. Chemicals used in this study.** Name, abbreviation where applicable, catalog number and company are provided for each chemical.

| **Primary antibody** | **Reducing agent** | **Blocking solution** | **Dilution** | **Company** | **Catalog number** |
| --- | --- | --- | --- | --- | --- |
| **AMPKα** | **TCEP** | **5% BSA** | **1:500** | **Cell Signaling Technology** | **5832** |
| **pAMPKα (Thr172)** | **TCEP** | **5% BSA** | **1:500** | **Cell Signaling Technology** | **2531S** |
| **ACC1** | **TCEP** | **5% BSA** | **1:1000** | **Proteintech** | **21923-1-AP** |
| **pACC1 (Ser79)** | **TCEP** | **5% BSA** | **1:1000** | **Proteintech** | **29119-1-AP** |
| **CPT1B** | **DTT** | **5% Milk** | **1:500** | **Proteintech** | **22170-1-AP** |
| **CPT1A** | **DTT** | **5% Milk** | **1:4000** | **Proteintech** | **15184-1-AP** |
| **CPT2** | **TCEP/DTT** | **5% Milk/5% BSA** | **1:2000** | **Abcam** | **AB151114** |
| **H6PD** | **DTT/TCEP** | **5% Milk** | **1:1000** | **Sigma-Aldrich** | **HPA004824** |
| **β-Tubulin** | **DTT/TCEP** | **5% Milk/5% BSA** | **1:3000** | **Proteintech** | **10094-1-AP** |
| **β-Actin** | **DTT/TCEP** | **5% Milk/5% BSA** | **1:8000** | **Santa Cruz** | **sc-47778** |

**Supplemental Table S2. Antibodies used in this study.** Sample reducing agent, membrane blocking solution, antibody dilution (in indicated blocking solution), antibody provider and corresponding catalog number are provided for each antibody used for immunoblotting.

| Gene | Primer sequences (forward, reverse) | Run method |
| --- | --- | --- |
| *18S* | AGTCCCTGCCCTTTGTACACA,  CGATCCGAGGGCCTCACTA | 95°C 5 min  PCR (40 cycles):  95°C 10 s, 60°C 15 s, 72°C 20 s  Melt curve:  95°C 15s , 60°C 60 s, 95°C 15 s |
| *Prkaa1* | CCTGGAGAAAGATGGCGAC, TCACAGCCACTTTATGTCCG |  |
| *Stk11* | CGAGGGATGTTGGAGTATGAG,  CTTAGTGTCTGGGCTTGGTG |  |
| *Camkk2* | AAGGGCCTGTAATGGAAGTG,  GGAGGTTGGAAGGTTTGATGTC |  |
| *Srebf1* | CCATCGACTACATCCGCTTC,  GCCCTCCATAGACACATCTG |  |
| *Ppara* | TGCAAACTTGGACTTGAACG, GATCAGCATCCCGTCTTTGT |  |
| *Ppargc1a* | AATGCAGCGGTCTTAGCACT,  ACGTCTTTGTGGCTTTTGCT |  |
| *Ppargc1b* | TGCGGAGACACAGATGAAGA,  GGCTTGTATGGAGGTGTGGT |  |

**Supplemental Table S3. Primer sequences and RT-qPCR run method used in this study.** Target gene name, forward and reverse primer sequences (5'-3') and corresponding qPCR cycling conditions are provided.

**Supplemental Table S4. Proteome-wide comparison between H6PD-/- and H6PD+/+ liver samples.** Quantitative proteomics results showing all detected proteins and their relative abundance changes between H6PD-/- and H6PD+/+ samples in liver homogenates and microsomal fractions. The table is provided as an Excel file in the Supplementary information.
